# An interneuronal CRH and CRHBP circuit stabilizes birdsong performance

**DOI:** 10.1101/2025.08.16.670679

**Authors:** Bradley M. Colquitt, Michael S. Brainard

## Abstract

The performance of skilled behaviors requires a balance between consistency and adaptability. Although the neural mechanisms that regulate this balance have been extensively studied at systems and physiological levels, relatively little is known about how the molecular properties of motor circuits influence motor stability versus flexibility. Here, we characterize the region- and cell-type specific expression patterns of neuropeptide systems across the neural circuit that controls the learning and performance of birdsong, a model for skilled behavior. We identify a number of neuropeptide pathways with differential expression between song regions and surrounding areas that are not involved in song production or learning. One of the strongest enriched genes in song regions is corticotropin releasing hormone binding protein (CRHBP), whose product binds corticotropin releasing hormone (CRH), a neuropeptide implicated in neuronal excitability and plasticity. We find that the expression of CRHBP in the song motor pathway decreases upon deafening-induced song destabilization, increases during song acquisition, and increases the more a bird sings. CRH and CRHBP are expressed in distinct interneuronal populations in song motor regions, providing a local neuromodulatory circuit well-positioned to regulate song performance. Consistent with this role, genetic and pharmacological manipulation of the CRH pathway in the song motor pathway resulted in bidirectional modifications of song variability, with elevated CRHBP acting to maintain low variability and elevated CRH acting to increase variability. These data indicate that an interneuronal neuropeptidergic pathway maintains the stability of song, acting as a local mechanism that regulates the balance between motor consistency versus flexibility.

## Introduction

Motor variability is actively regulated by motor circuits and is not simply a product of noise in the nervous system (Kao et al., 2005; Sober et al., 2008; Dhawale et al., 2017; Wu et al., 2014). A hallmark of successful skill learning is the gradual reduction of performance variability as actions become more precise. Reaching, kicking, and stepping begin as highly variable motor behaviors in infants that stabilize during childhood to become stereotyped actions (Adolph et al., 2018). Similarly, our first attempts at speaking take the form of acoustically variable vocalizations (“babbling”) that gradually transform into consistently performed phonemes. Motor variability has been extensively studied in terms of changes to neural connectivity (Aronov et al., 2008; Garst-Orozco et al., 2014), motor ensembles (Hinnekens et al., 2023; Peters et al., 2014), and dendritic spine dynamics (Chen et al., 2015; Roberts et al., 2010; Xu et al., 2009). However, we lack a clear definition of the molecular and cellular states of motor circuits that regulate performance variability.

The courtship song of songbirds is a complex, learned motor skill that serves as a powerful model for the neural mechanisms underlying motor learning and performance. Once learned, birdsong performance in many species is highly precise from rendition to rendition. However, the degree of this precision varies depending on social context, time-of-day, and the recency of song performance, indicating that it is an actively regulated motor property (Q. Chen et al., 2013; Hayase et al., 2018; Hilliard et al., 2012; Kao and Brainard, 2006; Kojima and Doupe, 2011; Miller et al., 2010; Ohgushi et al., 2015). Past work has identified systems-level circuit mechanisms that modulate song variability, most notably input from the song cortical-basal ganglia circuits to the song motor system (Andalman and Fee, 2009; Charlesworth et al., 2012, 2011; Kao et al., 2005; Olveczky et al., 2005; Warren et al., 2011). However, it remains relatively poorly understood how the local properties of the song motor circuit contribute to different aspects of song performance.

Neuropeptides play key roles in modulating the activity of neural circuits through their influence on synaptic transmission and the physiological properties of target neurons (Nusbaum and Blitz, 2012). Work in several model systems suggests that they are well-positioned to regulate motor output. In the crab stomatogastric ganglion, neuropeptide signaling modulates the functional connectivity between central pattern generators, resulting in different motor outputs depending on context (Dickinson et al., 1990). In nematodes, neuropeptides influence the activity of motor neurons to regulate basal locomotion and food-searching behavior (Buntschuh et al., 2018; Ramachandran et al., 2021). However, it is largely unknown how neuropeptides influence the performance of skilled behaviors in vertebrates, including the regulation of motor variability and the ability of a behavior to undergo adaptive plasticity.

To better understand the roles of neuropeptides on motor skill performance, we analyzed the expression of neuropeptides, their receptors, and regulatory factors across the neural circuit that controls birdsong as well as several comparator regions that have connectivity and physiology similar to the song circuit but that do not control song. We further analyzed how neuropeptide-associated gene expression changes during deafening-induced song destabilization (a process that results in increased song plasticity and variability), during normal song acquisition in juveniles (characterized by the reduction of plasticity and variability), and across varying rates of song performance (also linked to changes in song variability). We find that expression of corticotropin releasing hormone binding protein, CRHBP, is dynamically regulated during song destabilization, during song acquisition, and across variation in the number of songs a bird sings in a day. To test the role of the CRH system in regulating song performance, we performed targeted knockdowns and pharmacological manipulations of the CRH pathway and found that CRH and CRHBP have opposing effects on song variability. Together, these results highlight a role for neuropeptide systems in the regulation of motor performance through local regulation of central motor circuits.

## Results

### Expression of neuropeptide signaling systems in the song system

Birdsong is controlled by a dedicated neural circuit called the “song system” that allows for precisely interrogating how the molecular and cellular properties of a given song region influence song performance and learning (Fig. 1A,B). The song system comprises three cortical-like, or “pallial”, regions — HVC (acronym used as a proper noun), RA (robust nucleus of the arcopallium), LMAN (lateral magnocellular nucleus of the anterior nidopallium) — and one region in the basal ganglia, Area X. HVC and RA comprise the song motor pathway (SMP) and are necessary for song performance (Nottebohm et al., 1976). HVC influences the timing and temporal structure of song and projects to RA, which provides descending motor control of song via projections to syringeal and respiratory brainstem regions (Wild, 1993). The molecular, physiological, and connectivity properties of RA and HVC are highly similar to cortical extratelencephalic layer 5 and intratelencephalic neurons in mammals, respectively, and are functionally analogous to mammalian motor cortical circuits (Colquitt et al., 2021; Jarvis, 2019; Pfenning et al., 2014). Each song region is embedded in a region that has similar molecular and physiological properties but that do not directly influence song performance or learning: RA is located in the arcopallium (Arco), HVC in the caudal nidopallium (NC), LMAN in the rostral nidopallium (NR), and Area X in the striatum (Stri). These ‘non-song’ regions serve as comparators that enable the identification of gene expression profiles associated with song-dedicated regions.

**Figure 1.**
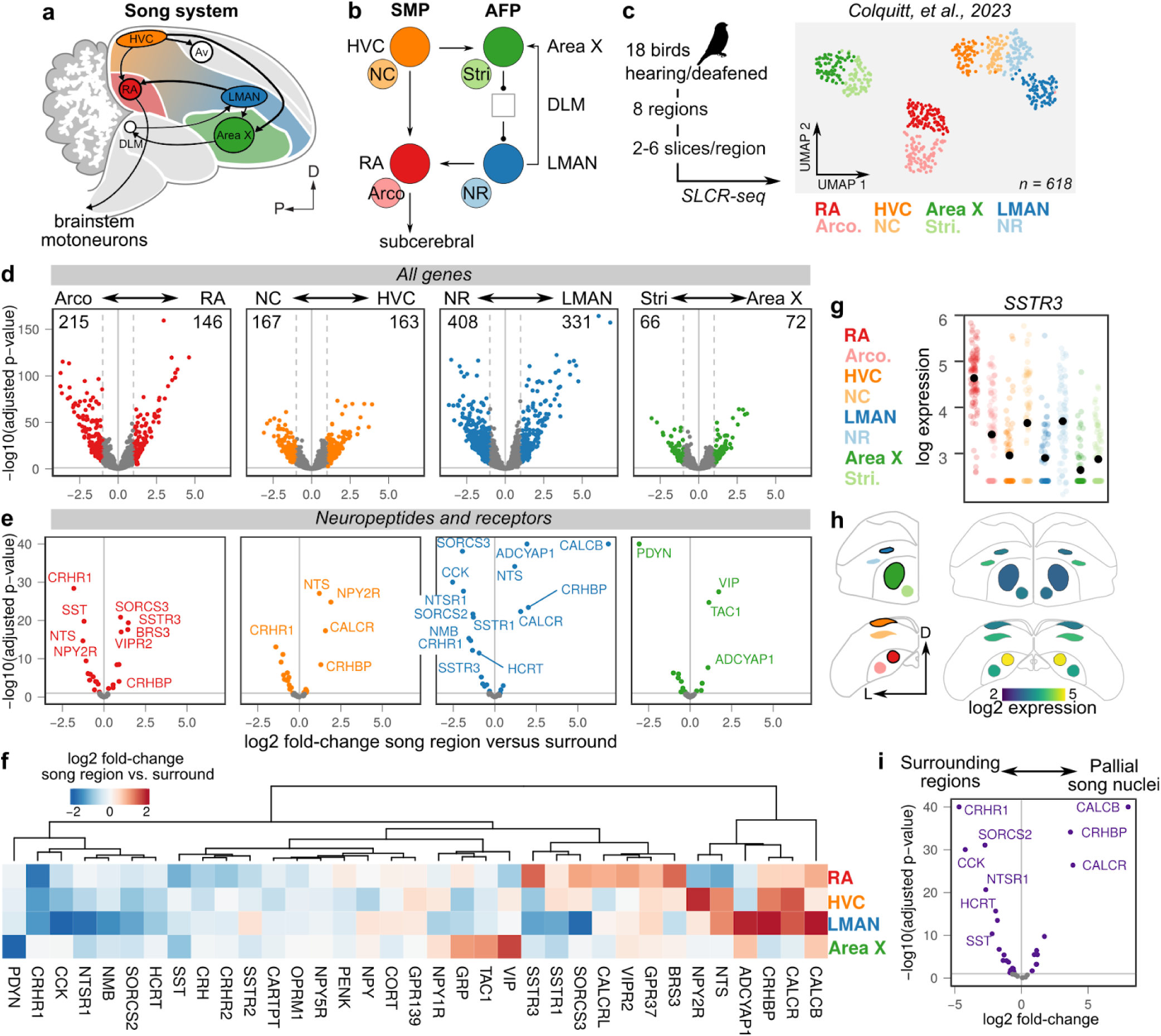
Specialized neuropeptide pathway expression in the song system. **(a)** Schematic overview of the song system. HVC, proper name; RA, robust nucleus of the arcopallium; LMAN, lateral magnocellular nucleus of the nidopallium; Av, avalanche; DLM, medial portion of the dorsolateral thalamic nucleus; D, dorsal; P, posterior. **(b)** Circuit diagram of the song system. Arrowheads and closed circles indicate excitatory and inhibitory connections, respectively. NC, caudal nidopallium; Arco., arcopallium; NR, rostral nidopallium; Stri., striatum. **(c)** Left: Experimental overview of SLCR-seq on hearing and deaf birds, from Colquitt et al. 2023. After a baseline period of song recording, birds were either deafened through bilateral cochlear removal or underwent a sham surgery. After 4, 9, or 14 days post-surgery, birds were euthanized and SLCR-seq libraries were prepared from HVC, NC, RA, Arco., LMAN, NR, Area X, and Stri. Right: Uniform Manifold Approximation and Projection (UMAP) plot of SLCR-seq data colored by section position. Each point reflects the gene expression profile of a single SLCR-seq sample. Samples show segregation by broad anatomical area — striatal (Area X), nidopallial (HVC, NC, LMAN, NR), arcopallial (RA, Arco.) — and song system nuclei from surrounding areas. **(d)** Volcano plots showing differentially expressed genes between each song and paired non-song region across all genes. Numbers in the top left and right indicate the number of genes with adjusted p-values < 0.1 and an absolute log2 fold-change greater than 1. **(e)** Volcano plots as in (d) subsetted for neuropeptide-associated genes (ligands, receptors, and modulators; see Methods for definition). **(f)** Patterns of neuropeptide gene expression enrichment and depletion in song regions relative to surround regions. **(g)** Normalized log gene expression data of an example gene *SSTR3* whose expression is enriched in RA relative to Arco, and depleted in HVC and LMAN relative to their surrounding regions. Each point indicates gene expression in a single SLCR-seq sample. **(h)** Coronal anatomical atlas representation of the expression of *SSTR3*. Left: Each region shown with color scheme as in A. Right: log gene expression value. D, dorsal; L, lateral. **(i)** Analysis of differential expression between the three pallial (cortical) song nuclei, HVC, RA, and LMAN and their surrounding regions, NC, Arco, and NR.

In a recent study, we generated a set of bulk RNA-seq libraries encompassing RA, HVC, LMAN, Area X, and matched non-song regions for each song area, yielding eight regions total (Fig. 1C) (Colquitt et al., 2023). These libraries were generated using an approach we designed called Serial Laser Capture RNA (SLCR)-seq, which allowed the collection of multiple (2 to 6) thin cryosections from each brain region from each bird (see Methods). Individual RNA-seq libraries were then generated from each cryosection. This approach was applied to 18 adult Bengalese finch males, nine of which had been deafened through bilateral cochlear removal and the remaining nine of which had undergone a sham surgery. In total, this dataset contains 618 individual SLCR-seq libraries.

To identify neuropeptide systems that are differentially expressed between each song region and its complementary non-song comparator region (e.g. RA vs. Arco and HVC vs. NC), we analyzed differential expression in the SLCR-seq dataset across all genes (Fig. 1D, see Methods for analysis). For each comparison, we identified a large set of differentially expressed genes (138-739, adjusted p-value < 0.1, absolute log fold-change > 1), consistent with previously identified transcriptional specializations of the song system (Lovell et al., 2013, 2008; Mello et al., 2019; Nevue et al., 2020).

We subsetted these genes for neuropeptides, neuropeptide receptors, and neuropeptide modulators (see Methods for list definition), yielding 80 genes in total. Each song region exhibited differentially expressed neuropeptide-related genes relative to surround (16 to 23 genes) (Fig. 1E). Diverse patterns of neuropeptide-related gene expression across song regions suggest that each region has a distinct neuromodulatory state (Fig. 1F-H and Fig. S1). For example, somatostatin receptor 3 (*SSTR3*) is strongly enriched in RA but relatively depleted in other song regions (Fig. 1G and H). Similarly, neuropeptide Y receptor Y2 (*NPY2R)* is strongly enriched in HVC but no other song region (Fig. S1), and vasoactive intestinal peptide (*VIP)* is strongly elevated in the striatal song region Area X (Fig. S1).

We also determined which genes have expression that is significantly enriched or depleted in the pallial (cortical) song regions, RA, HVC, and LMAN, as a group relative to surrounding non-song regions (Fig. 1I). Calcitonin related polypeptide beta (*CALCB*) and its receptor *CALCR* are enriched in each of the pallial song regions, with particularly strong enrichment seen in LMAN (Fig. 1F and I). Likewise, corticotropin-releasing hormone binding protein (*CRHBP*) is enriched in each of the pallial song regions while the CRH receptor *CRHR1* is depleted relative to surrounding areas. The expression of two neuropeptides cholecystokinin (*CCK*) and *SST* are depleted across RA, HVC, and LMAN.

In summary, each song region shows distinct specializations of both neuropeptide ligand and receptor expression relative to surrounding non-song regions.

### Patterns of neuropeptide signaling-system expression across cell types in the song system

Neuropeptide signaling systems form local modulatory networks across cell types. To examine the distributions of neuropeptide ligands and receptors across cell types in the song system, we analyzed a single-cell and single-nucleus dataset from zebra finches and Bengalese finches(Colquitt et al., 2021) (Fig. 2A and B). This dataset was generated from the two regions of the song motor pathway, HVC and RA, and consists of a range of glutamatergic and GABAergic neuron types, their neurogenic precursors, and non-neuronal cells (astrocytes, oligodendrocytes, oligodendrocyte precursor cells (OPCs), microglial, endothelial cells, and red blood cells). HVC contains five glutamatergic cell clusters with HVC_Glut-1 and HVC_Glut-3 corresponding to HVC_RA_ and HVC_X_ projection neurons, respectively (Fig. 2C). RA glutamatergic neurons project to subcerebral targets, and the three transcriptionally defined clusters in this dataset have distinct but overlapping projection patterns (Colquitt et al., 2021). GABAergic neuron clusters are well-conserved between birds and mammals and can be clustered into lateral, medial, and caudal ganglionic eminence (L/M/CGE) types. GABA-1 neurons are strongly similar to LGE-type neurons from the mammalian amygdala and olfactory bulb; GABA-2/3/4 clusters form a spectrum between SST and PVALB-class MGE neuron types as defined for the mammalian cortex; GABA-5 neurons are strongly similar to Vip/Sncg class neurons from the CGE; GABA-6 neurons are similar to Lamp5-class neurogliaform neurons; GABA-7 neurons have an MGE-like transcriptional signature but do not cleanly map onto a mammalian neuron type; and GABA-8 neurons are similar to CGE cholinergic neurons.

**Figure 2.**
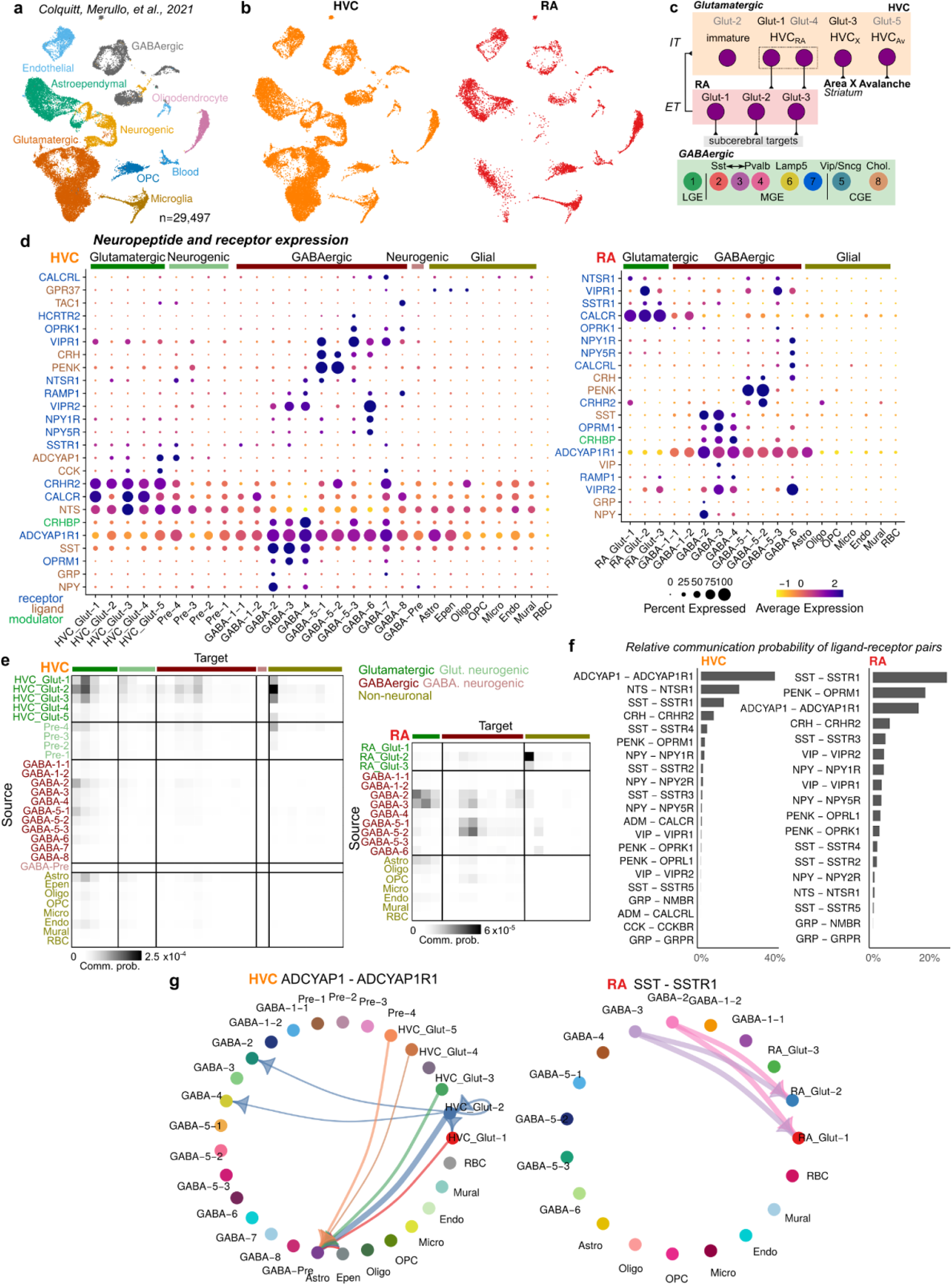
Patterns of cell type specific expression of neuropeptide systems in the song motor pathway. **(a,b)** Uniform manifold approximate projection plots of single-nucleus RNA-sequencing from two song regions (RA and HVC) adapted from (Colquitt et al., 2021). (a) Points colored and labeled by cluster and (b) points colored and split by region. **(c)** Schematic of identified clusters in each region. GABAergic neuron naming convention follows that used in (Colquitt et al., 2021). **(d)** Expression of neuropeptide ligands and receptors across HVC and RA cell types. **(e)** CellChat analysis of overall neuropeptide cell-cell communication probability in HVC and RA. **(f)** Predicted relative communication probability of neuropeptide ligand-receptor pairs in each region (see Methods). Communication probabilities were summed across cell types for each ligand-receptor pair, then normalized by the total communication probability. **(g)** Communication probabilities for ADCYAP1-ADCYAP1R1 and SST-SSTR1, the top two ligand-receptor pairs in HVC and RA, respectively. Arrow widths are scaled to communication probabilities. The top 1% of interactions are shown.

Neuropeptides and their receptors displayed diverse expression patterns across these cell types (Fig. 2D). GABAergic neuron types in particular showed specific expression of a number of different neuropeptides and their receptors, suggesting dense intercellular neuropeptidergic modulation among inhibitory interneurons in the song motor pathway. In both HVC and RA, glutamatergic neurons express fewer neuropeptide ligands and receptors, but do show moderate to strong expression of a handful of receptors, including *CALCR*, *CRHR2, VIPR1/2, NTSR1,* and *SSTR1*, suggesting they are a capable of responding to diverse signaling systems. Non-neuronal cell types largely did not express neuropeptide components with the notable exception of astrocytes, which exhibited strong expression of *ADCYAP1R1* (Adenylate Cyclase Activating Polypeptide 1 (Pituitary) Receptor Type I), a member of the pituitary adenylate cyclase-activating peptide (PACAP) ligand/type 1 receptor (PAC1) system. Previous work indicates that this system regulates glutamate release from astrocytes in the mammalian neocortex (Kong et al., 2016), pointing to an underappreciated role for neuron-astrocyte communication in birdsong motor circuit function.

To quantify the patterns of intercellular communication across cell types, we analyzed these datasets using CellChat (Jin et al., 2021, 2023), an approach that models between-cell communication probability according to known interactions among ligands and receptors, the expression level of these pairs across cell types, and the relative abundance of each cell type (Fig. 2E-G). An analysis of overall communication between cell types across all neuropeptide systems indicates that several GABAergic cell types are prominent senders (ligand-expressing) and receivers (receptor-expressing) in both RA and HVC (Fig. 2E). Notably, however, communication among GABAergic cell types is low in HVC relative to RA. In HVC, glutamatergic neurons are prominent neuropeptide sources, targeting receptors in both glutamatergic and GABAergic neurons. In contrast, RA glutamatergic neurons are relatively weak neuropeptide sources. One notable exception to this pattern is for astrocytes, which are prominent targets of neuropeptides expressed in both HVC and RA glutamatergic neurons.

We next analyzed which ligand-receptor pairs display the strongest communication probability in each song nucleus (Fig. 2F and G and Fig. S2-S4). HVC and RA employ similar signaling systems with some distinct differences. ADCYAP1-ADCYAP1R1, SST-SSTR1, and CRH-CRHR2 are all within the top 5 pairs for each song region. Interestingly, in both regions ADCYAP1-ADCYAP1R1 appears to be the sole signaling system driving the observed interaction between glutamatergic neurons and astrocytes (Fig. 2G and Fig. S2-S4). SST-SSTR1 (and to a lesser extent SSTR3 and SSTR4) form a prominent connection from GABA-2/3/4 (MGE *Sst*/*Pvalb*-class) neurons to glutamatergic neurons in HVC and RA (Fig. 2G, S3, and S4). Similarly, in both regions CRH-CRHR2 signaling spans from GABA-5 (CGE *Vip*/*Sncg*-class) neurons to glutamatergic neurons (Fig. S2 and S3). Opioid signaling, here represented by PENK-OPRM1, shows strong interneuron-interneuron communication from GABA-5 to GABA-2/3/4 in both regions as well as GABA-5 to glutamatergic neurons in HVC. In contrast, to mammalian cortical systems, VIP is not expressed in “*Vip-*class” interneurons (GABA-5) in HVC and RA (Fig. 2D and Fig. S3,S4). Rather, it is predominantly expressed in GABA-3 interneurons, a *Sst/Pvalb*-class neuron type, albeit at low levels in HVC. The VIP receptors *VIPR1* and *VIPR2* are expressed across multiple neuron types, suggesting broad influence of the neuropeptide across the local microcircuits of the song motor pathway.

**Figure 3.**
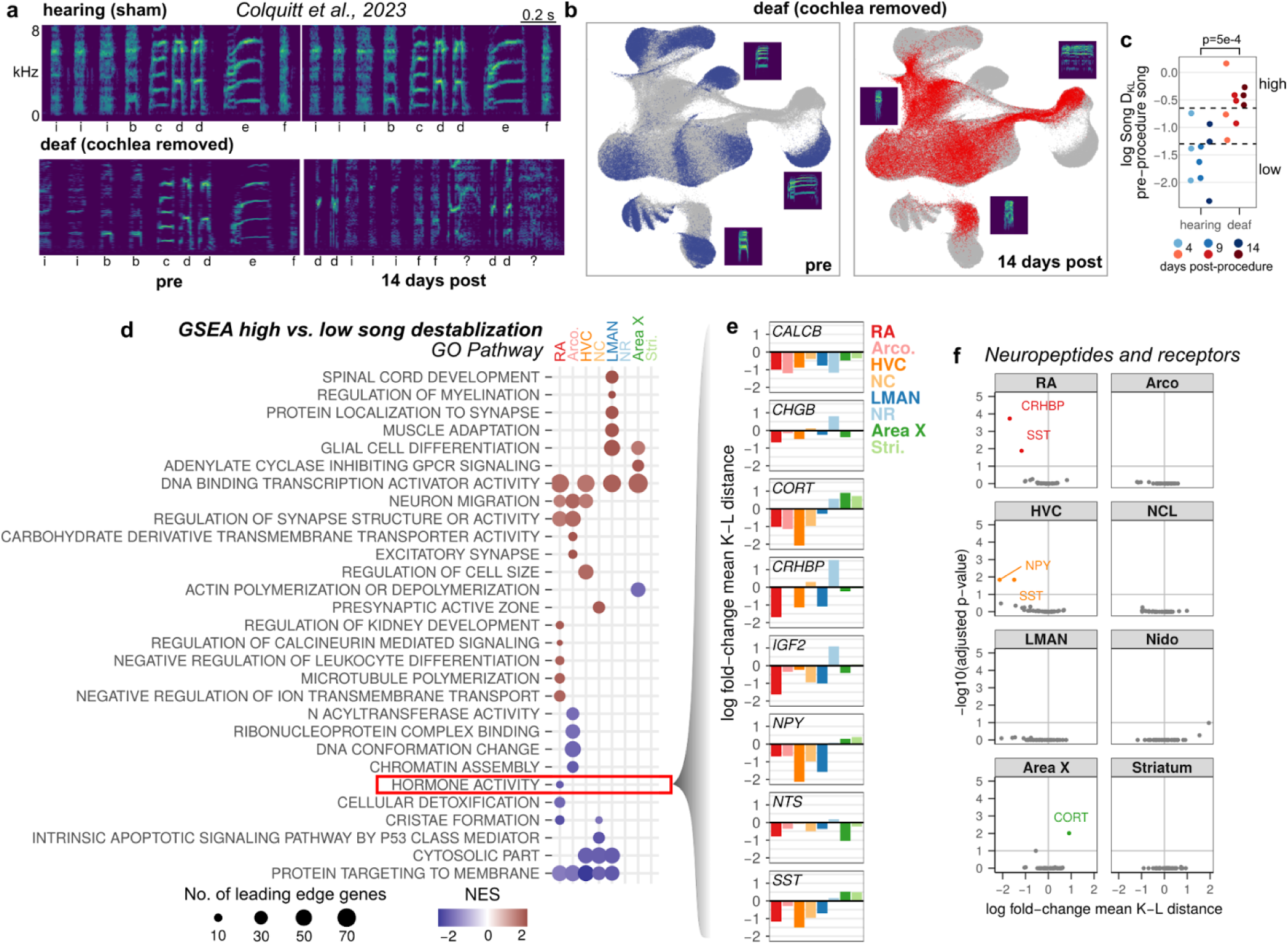
Song destabilization results in reduced expression of several neuropeptide-associated genes in the motor output regions RA and HVC. **(a)** Example spectrograms from one hearing (sham) and one matched (sibling) deaf (bilateral cochlea removal) bird. Songs are shown from before the procedure and 14 days following the procedure. Labels below each spectrogram correspond to discrete categories of song units (‘syllables’). kHz, kiloHertz. **(b)** Uniform Manifold Approximation and Projection (UMAP) representation of syllable spectrograms (see Methods) across the entire recording period for the deafened bird (4 days before to 14 days after the procedure). Data are split into ‘pre’-procedure (blue: 4 to 1 day before surgery) and ‘post’-procedure (red: 14 days after surgery) subsets. For reference, gray points in each plot correspond to data across the entire recording period. Example syllable spectrograms are placed adjacent to their position in UMAP space. **(c)** Relative spectral distance between syllables pre- and post-procedure, represented as the mean Kullback-Leibler (KL) distance between Gaussian mixture models or ‘Song D_KL_’ (see Methods for calculation). Song DKL trends higher with increasing days from deafening. Significance calculated using a two-sided Wilcoxon rank-sum test. **(d)** Gene set enrichment analysis (GSEA) of song destabilization-associated genes. Shown are the Gene Ontology (GO) terms that are significant in at least one song or non-song region (adjusted p-value < 0.1, see Methods). Dot color indicates the normalized enrichment score, and dot size indicates the number of leading edge genes (core genes most responsible for the enrichment associated with the GO term). Terms are ordered by hierarchical clustering (Euclidean distance, Ward squared method). Highlighted is the term Hormone Activity (GO:0005179). **(e)** Log2 fold-change expression of the leading edge genes in term Hormone Activity across song and surround regions. **(f)** Volcano plot of destabilization-associated differential expression, subsetted for neuropeptide-associated genes.

### Song destabilization after deafening results in a reduction of neuromodulatory genes

The range of expression patterns of neuropeptide ligand and receptor across cell types in the song motor pathway suggests that these signaling systems may regulate birdsong performance and plasticity. Deafening through cochlea removal destabilizes birdsong over the course of several days, rendering it both less structured and more variable from rendition-to-rendition (Fig. 3A,B) (Nordeen and Nordeen, 1992; Okanoya and Yamaguchi, 1997; Woolley and Rubel, 1997; Brainard and Doupe, 2001; Colquitt et al., 2023). In our previous study, we identified patterns of gene expression perturbations across the song system that correlate with deafening-induced birdsong destabilization, represented by a measure called Song D_KL_ that provides an aggregate statistic of how much a bird’s song changes after cochlea removal (Fig. 3C). Gene set enrichment analysis indicated that a number of biological processes exhibit coherent up- or down-regulation with song destabilization in the various regions of the song system. In particular, genes associated with hormone activity (GO:0005179) showed a significant reduction in expression in RA with higher degrees of song destabilization (Fig. 3D). The leading edge genes in this set comprise a number of neuropeptides (*SST, NPY, CORT, IGF2, NTS, CALCB*), neuropeptide modulators (*CRHBP*), and proteins involved in neuropeptide secretion (*CHGB*) (Fig. 3E). Similarly, the expression levels of *CRHBP* and *SST* were significantly reduced with song destabilization in RA, while *SST* and *NPY* levels were reduced in HVC (Fig. 3F). Adjacent non-song regions showed no significant alterations of neuropeptide-associated gene expression, indicating that this is a neural circuit-specific response.

### CRHBP expression increases in the song motor pathway during song acquisition

Across the signaling systems affected by deafening, we were especially interested in investigating links between corticotropin releasing hormone signaling pathways and song destabilization. In particular, prior work has established a role for CRH signaling in modulating neural excitability and plasticity (Gallagher et al., 2008). Consistent with its role in modulating synaptic and physiological properties, CRH has been shown to influence a number of neural processes, including learning and memory in the hippocampus (Chen et al., 2012) and anxiogenic behavior in the prefrontal cortex (Li et al., 2016).

CRHBP is a secreted glycoprotein that binds to corticotropin-releasing hormone (CRH, also known as corticotropin-releasing factor) (Kemp et al., 1998; Ketchesin et al., 2017). In most systems examined, CRHBP inhibits CRH signaling either by acting as a ligand sink or by influencing the kinetics of CRH binding to its receptors CRHR1 and CRHR2 (however, see (Ungless et al., 2003) for an example of a facilitating interaction between CRHBP and CRH). The reduction of *CRHBP* expression after the destabilization of adult birdsong suggests that the regulation of this gene is sensitive to the plasticity state of song and could play a role in deafening-induced song plasticity.

In contrast to the destabilization of song that occurs following deafening, song becomes more stable during its initial developmental learning. Male Bengalese finches learn their songs during the first few months of life in several stages (Fig. 4A). Song begins around the first month after hatching as a highly variable and poorly structured vocalization called ‘subsong’ (akin to babbling). It then progresses through a ‘plastic’ stage for several months (50-90 days post hatch, dph) in which its acoustic and temporal features become more structured yet remain variable. Finally, song enters a ‘crystallized’ stage (>90 dph) evidenced by a highly precise and well-structured vocalization.

**Figure 4.**
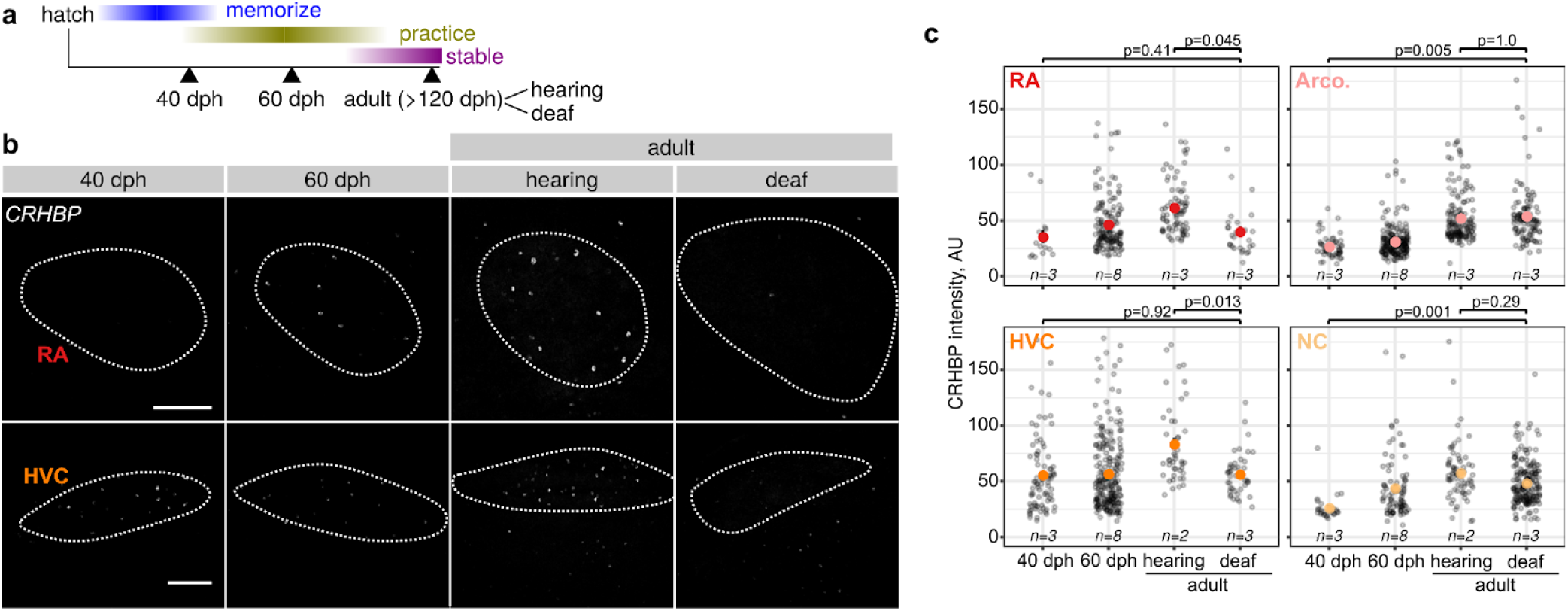
*CRHBP* expression in the song motor pathway increases during song acquisition and decreases following deafening. **(a)** Developmental timeline of birdsong learning. dph, days post hatch. **(b)** Representative images of *in situ* hybridizations against *CRHBP* in RA and HVC (outlined) from birds at different developmental stages or adult birds that are hearing or deaf. Scale bar is 200 μm. **(c)** Quantification of signal intensities across conditions and brain regions. Each point is the *CRHBP* signal intensity in each *CRHBP-*positive cell. P-values are derived from linear mixed models (see Methods).

To determine whether *CRHBP* levels correlate with developmental song stabilization, we examined *CRHBP* expression during initial song learning in juveniles using *in situ* hybridization on brain sections from birds in the subsong (40 dph), plastic (60 dph), and stable phases (‘adult’, 142-269 dph) as well as on sections from birds that had been deafened (Fig. 4B and C). The birds in the hearing and deafened conditions were independent from those used in the SLCR-seq dataset. *CRHBP* levels increased in RA and HVC as well as each surrounding region Arco. and NC from subsong to adult stages. However, following deafening, *CRHBP* levels dropped to subsong levels only in the song regions RA and HVC, consistent with the SLCR-seq results.

### CRHBP expression in the song motor pathway is correlated with singing

Singing influences gene expression in the song system, likely through gene regulatory mechanisms that are sensitive to neural activity (Q. Chen et al., 2013; Feenders et al., 2008; Hayase and Wada, 2018; Hilliard et al., 2012; Horita et al., 2012; Jarvis et al., 1998; Jin and Clayton, 1997; Kimpo and Doupe, 1997; Li et al., 2000; Poopatanapong et al., 2006; Sasaki et al., 2006; Wada et al., 2006; Warren et al., 2010; Whitney et al., 2014; Whitney and Johnson, 2005). In dissociated mouse cortical neurons, depolarization induces the expression of *CRHBP* and *SST* (Tyssowski et al., 2018) and *CRHBP* levels increase in the hippocampus following seizures (Smith et al., 1997). In addition, the amount a bird sings influences the variability of song, suggesting that activity-dependent mechanisms may influence song consistency (Miller et al., 2010; Q. Chen et al., 2013; Ohgushi et al., 2015; Hayase et al., 2018; Mizuguchi et al., 2024).

To determine whether singing modulates the expression of neuropeptides generally and the song stabilization-associated genes *CRHBP* and *SST* specifically in the song system, we regressed song system gene expression from the SLCR-seq dataset against two measures of singing amount: (1) the number of songs sung in the two hours before euthanasia (‘2 hours’) and (2) the number of songs sung from two days before euthanasia up to the time of euthanasia (‘2 days’) (Fig. 5A and B). These measures are correlated (Fig. S5A, Pearson *R* = 0.83, p = 1.9e-5) yet capture different aspects of song-related activity, the first reflecting singing differences more proximal to the time gene expression was profiled and the second representing longer-scale variation in singing amount. To identify coordinated responses that could reflect neural activity variation throughout the song system, we focused on genes that exhibited similar fold-changes across the three pallial regions of the song system, RA, HVC, and LMAN (33 genes total, adjusted p-value < 0.1, Fig. 5C and Fig. S5B). These song-modulated genes were enriched for several biological functions, notably cellular respiration, neuromodulation, neurotransmission, and regulation by neural activity (Fig. 5D). Among the neuromodulation-associated genes were several neuropeptide-associated genes, including *CRHBP*, *SST, NPY, CHGB,* and *SCG2.* Inspection of the extent to which the expression of *CRHBP* and *SST* expression varied with the amount of singing indicated that, although both showed some degree of variation, *CRHBP* exhibited greater dependency on singing amount than *SST* across the song system (Fig. 5E and Fig. S5C).

**Figure 5.**
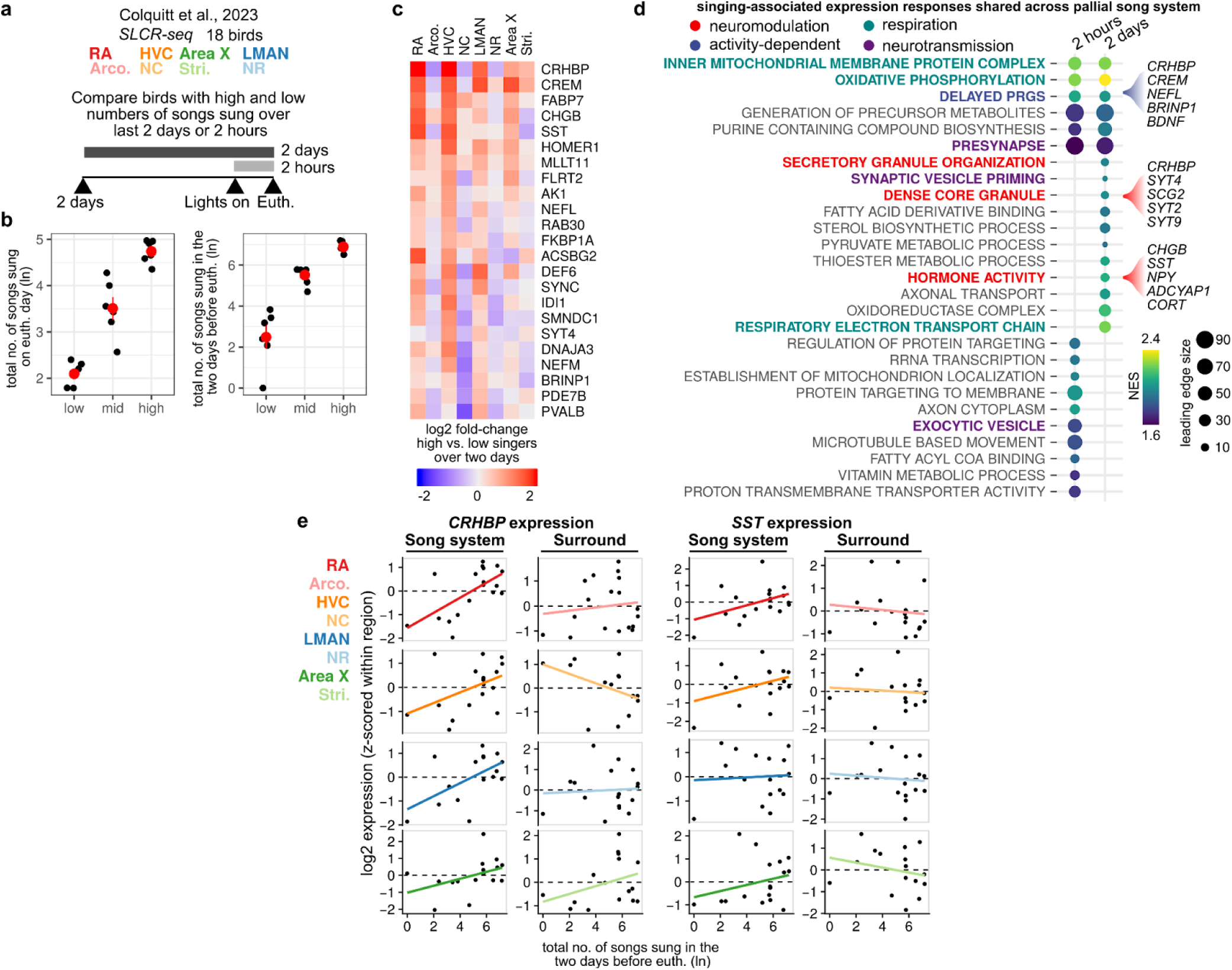
Modulation of neuropeptide system expression in the song system by singing. **(a)** Analysis of singing-dependent gene expression across two different time scales: total number of songs sung during the two hours before euthanasia and total number of songs sung during the two days before euthanasia. **(b)** Total number of songs sung by the birds in the SLCR-seq dataset in the two hours (*left*) or two days (*right)* before euthanasia. Birds are divided into low, mid, and high groups for differential gene expression analysis. **(c)** Heatmap of genes showing significantly increased expression (adjusted p<0.1) across pallial song nuclei (RA, HVC, and LMAN) between 2 day high and low singers. **(d)** Gene set enrichment analysis of singing-rate associated expression responses that are shared across the pallial song system (for both two-hours and two-days measures). Enriched are several groups of terms including those related to neuropeptides, activity-dependent gene regulation, neurotransmission, and cellular respiration. Listed are the five top leading edge genes for selected gene sets related to neuropeptides (“Hormone activity”, “Dense core granule”) and regulation by neural activity (“Delayed PRG (primary response genes)”). **(e)** Singing-modulated expression of *CRHBP* and *SST* across the song system and surrounding regions for the two-days measure.

To validate these findings, we recorded the songs of eight adult males and euthanized them nine to ten hours after lights on (Fig. S6A). This later euthanasia time point allowed birds to sing more relative to the birds analyzed in the deafening experiment, generating a larger range of singing bout numbers. We analyzed the expression of *CRHBP* and *SST* by reverse transcription-quantitative PCR in RA and arcopallium. *CRHBP* expression in RA was correlated with the number of songs sung on the day of euthanasia and the mean number of songs sung per day (adjusted R^2^ = 0.75, p = 0.003; adjusted R^2^ = 0.53, p = 0.04, respectively) but showed no association with singing in arcopallium (adjusted R^2^ = 0.02, p = 0.3; adjusted R^2^ = 0.11, p = 0.4) (Fig. S6B). In contrast, *SST* expression showed no significant association with singing in either region, consistent with the weaker relationship of *SST* expression with singing relative to *CRHBP* seen in the SLCR-seq data. Using *in situ* hybridization in a distinct set of six birds, we validated the positive correlation between *CRHBP* expression in RA and singing and found a similar correlation in HVC (Fig. S6C and D), consistent with the SCLR-seq data (Fig. 5C and E). These results indicate that the expression of the song stabilization-associated gene *CRHBP* increases as birds sing more songs.

### Components of the CRH signaling pathway are expressed in distinct neuronal populations of the song motor pathway, representing a conserved neuropeptidergic circuit

The strong reduction of *CRHBP* expression in RA during song destabilization induced by deafening, its increase during stabilization of song over the course of developmental song learning, and its association with singing amount indicate that CRHBP is dynamically modulated in the song system. These correlations across multiple song conditions raises the hypothesis that CRH signaling in RA influences song performance. To test this hypothesis, we first examined the cell type-specific expression of each CRH signaling component.*CRHBP* is broadly enriched in the pallial components of the song system (HVC, RA, and LMAN), showing 1.1-1.6 log-fold enrichment in each song nucleus relative to surrounding regions (Fig. 6A and Fig. S7A). In contrast, *CRH* expression is reduced in RA relative to adjacent arcopallium and has moderate expression in other song and non-song regions. The CRH receptor *CRHR1* is expressed at low levels across the areas assayed and shows significant reduction in pallial song regions. *CRHR2* is more highly expressed and at similar levels between song and non-song regions.

**Figure 6.**
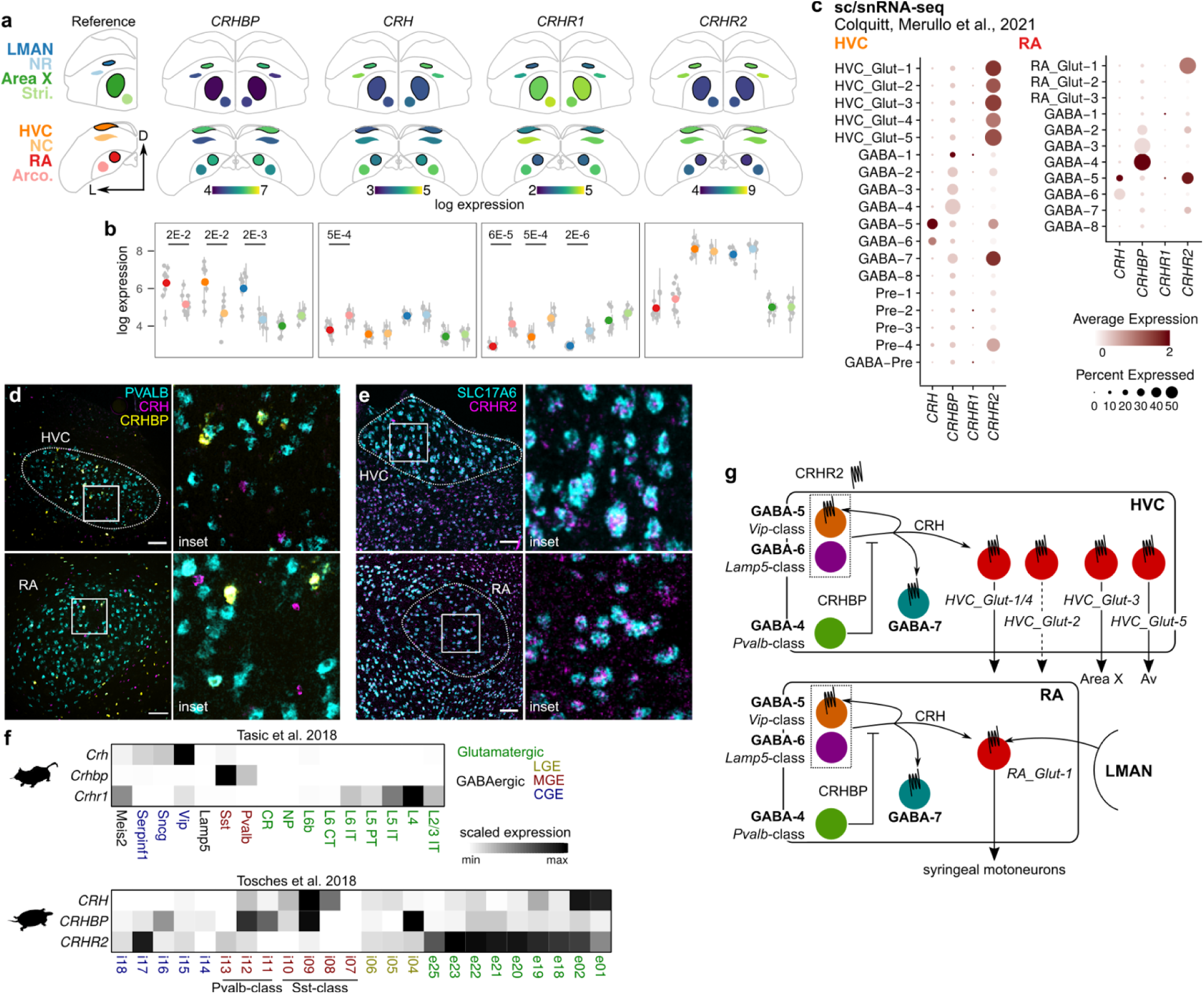
Distributed expression of the CRH neuromodulatory pathway in the song motor pathway. **(a, b)** Expression of CRH pathway components, *CRHBP, CRH*, *CRHR1*, *CRHR2*, in the song system and non-song surround regions. **(a)** Atlas representation of SLCR-seq data showing expression in coronal brain sections. **(b)** Point representation of SLCR-seq data. Each point is the average gene expression in a given region in a single bird. Error bars are standard error of the mean across 2-6 sections/region/bird. Colored dots represent mean expression across birds for a given region. Mean differences were calculated between each song and adjacent non-song region, and adjusted p-values (Student’s t-test, Benjamini-Hochberg adjustment) are shown for comparison with p < 0.05. **(c)** Expression of CRH pathway components in HVC and RA cell types. A previously generated single-cell/single-nucleus RNA-seq (scRNA/snRNA-seq) dataset (Colquitt et al., 2021) was queried for *CRH*, *CRHBP*, *CRHR1*, and *CRHR2* expression, represented here as a dotplot. Dot intensity reflects the average expression of a given gene in a given cluster (cell type), and dot size indicates the percentage of cells in a cluster that have detectable expression of a given gene. **(d, e)** *In situ* hybridization (ISH) validation of *CRH, CRHBP*, and *CRHR2* expression in the song system. **(d)** *CRH* and *CRHBP* show non-overlapping expression patterns in HVC and RA. *PVALB* is used here to demarcate HVC and RA which show elevated levels of *PVALB* in glutamatergic neurons (Wild et al., 2001). *PVALB* is also a general marker for GABA-2/3/4 (*Sst*-class and *Pvalb*-class) interneuron subtypes. **(e)** ISHs of the CRH receptor *CRHR2* in HVC and RA indicate broad expression in glutamatergic neurons (indicated by *SLC17A6*). Scale bar is 100 μm. **(f)** Expression of CRH pathway components *CRHBP*, *CRH, CRHR1*, and *CRHR2* in neurons of the mouse neocortex (Tasic et al., 2018) and turtle pallium (Tosches et al., 2018). Expression is scaled to the minimum and maximum of each gene. Cell cluster annotations are labels from the original publications. *Crhr2* in mouse and *CRHR1* are not shown due to low expression. **(g)** Model of the local CRH neuromodulatory pathway in HVC and RA. Arrows indicate positive interactions, blocked lines indicate negative interactions. CRH, released by *Vip-*class GABA-5 (and to a lesser extent *Lamp5*-class GABA-6), activates CRHR2 receptors expressed by glutamatergic neurons, GABA-7 interneurons, and GABA-5 interneurons (autoregulatory). CRHBP is released from GABA-4 *Pvalb-*class interneurons and binds to CRH, thereby reducing activation of CRHR2.

To examine the cell type expression of these components, we interrogated the previously published single-cell/nucleus RNA-sequencing dataset (Colquitt et al., 2021) obtained from microdissected HVC and RA, as described above. Each CRH pathway component shows similar cell type-specific expression patterns in both HVC and RA (Fig. 6C). *CRH* and *CRHBP* expression is restricted to different populations of GABAergic interneurons: *CRH* is most strongly expressed in GABA-5 interneurons (homologous to *Vip* and *Sncg*-class disinhibitory interneurons in mammals (Colquitt et al., 2021; Tasic et al., 2018) with lower expression in GABA-6 (homologous to *Lamp5-*class neurogliaform interneurons). *CRHBP* is enriched in GABA-4 interneurons, which show strong transcriptional similarity to mammalian fast-spiking *Pvalb*-class interneurons, as well as moderate expression in GABA-2 and GABA-3, which are similar to *Sst-*class interneurons. In contrast, the receptor *CRHR2* is expressed primarily in glutamatergic projection neurons and GABA-5 (the predominant *CRH*-expressing class). *In situ* hybridizations confirmed these patterns of expression, with *CRH* and *CRHBP* showing non-overlapping cell-type specific expression and broad expression of *CRHR2* in glutamatergic neurons (Fig. 6D and E).

Corticotropin releasing hormone signaling has been implicated in the modulation of forebrain neural circuits in mammals (Aldenhoff et al., 1983; Kratzer et al., 2013; Li et al., 2016), suggesting that this pathway might play similar roles across species. To determine if the different components of this pathway share a similar cellular expression pattern across amniotes, we examined the expression of *CRH*, *CRHBP*, and *CRHR1/2* in publicly available mouse cortical and turtle pallial single-cell RNA-sequencing datasets (Tasic et al., 2018; Tosches et al., 2018) (Fig. 6F). In both species, the cellular distribution of *CRHBP* and *CRHR1/2* was similar to that found in songbirds: *CRHBP* is enriched in *Sst/Pvalb-*class GABAergic neurons while the receptors (*CRHR1* in mouse and *CRHR2* in turtles) are predominantly expressed in glutamatergic neurons. *CRH* is expressed in *Vip*-class neurons in mouse neocortex, similar to songbirds, yet is expressed in a range of neuronal types in turtles. These similar patterns of expression suggest that the cellular distribution of the CRH pathway is conserved across amniotes.

In aggregate, these patterns of expression indicate that the different components of the CRH pathway are present in each song motor pathway nucleus and are distributed across different neuronal populations. This configuration supports a neuropeptidergic circuit model (Fig. 6G): CRHBP, released by fast-spiking interneurons (GABA-4), inhibits the ability of CRH, released from disinhibitory interneurons (GABA-5) and to a lesser extent from neurogliaform interneurons (GABA-6), to bind to CRH receptors expressed by glutamatergic projection neurons.

### Modulation of CRH and CRHBP bidirectionally affects song variability

This neuropeptidergic circuit model predicts that CRH and CRHBP would have opposing effects on song stability, due to the inhibitory influence of CRHBP on CRH. In particular, we predicted that reducing *CRHBP* expression, as was seen with increased levels of song destabilization, would release CRH from inhibition and result in increased song variability. Similarly, we predicted that increasing CRH levels would overcome CRHBP inhibition and result in increased song variability while reducing CRH levels would result in reduced song variability. To test these predictions, we performed two sets of manipulations — (1) expression knockdowns of CRHBP and CRH and (2) pharmacological activation of the CRH pathway — and examined their effects on song.

We first screened candidate siRNAs in primary neural cultures from Bengalese finch brains and selected 3 siRNAs targeting each gene that produced strong knockdown relative to transfections of a control siRNA (percent knockdown of expression relative to control: CRHBP 48-80%, CRH 58-91%, Fig. S8A). We then optimized an *in vivo* siRNA delivery method using a cationic lipid transfection reagent to knockdown *CRHBP* and *CRH* in specific stereotactically identified brain regions (see Methods). In validation assays using qRT-PCR, this approach produced at least ∼50% knockdown of targeted genes *in vivo* at two days after injection relative to paired control siRNA injections (Fig. S8B,C).

We stereotactically injected *CRHBP, CRH,* and control siRNAs bilaterally into RA of adult male birds and monitored effects on song (Fig. 7A). As with the analysis of song in hearing and deafened birds (Colquitt et al., 2023), we examined both global perturbations to song using dimensionality reduction of syllable spectrograms, as well as more targeted analyses of alterations to syllable fundamental frequencies. Knockdown of either *CRHBP* and *CRH* altered global song quality (Fig. S8D and E), with knockdowns of *CRHBP* having a greater effect than that of *CRH*. To further characterize the degree to which knockdowns altered song structure, we calculated the distance between syllables spectrograms from song before and after siRNA injections and compared these distances to those following deafening-induced song destabilization (Fig. 7B and C). Two to three days following injection, *CRHBP* knockdown disrupted syllable spectrograms to a similar level as destabilization following deafening, while *CRH* knockdown elicited a small but insignificant change relative to controls. These effects indicate that a causal reduction of *CRHBP* expression in RA of hearing birds, parallel to the reduction observed following deafening, is sufficient to mimic aspects of deafening-induced song destabilization.

**Figure 7.**
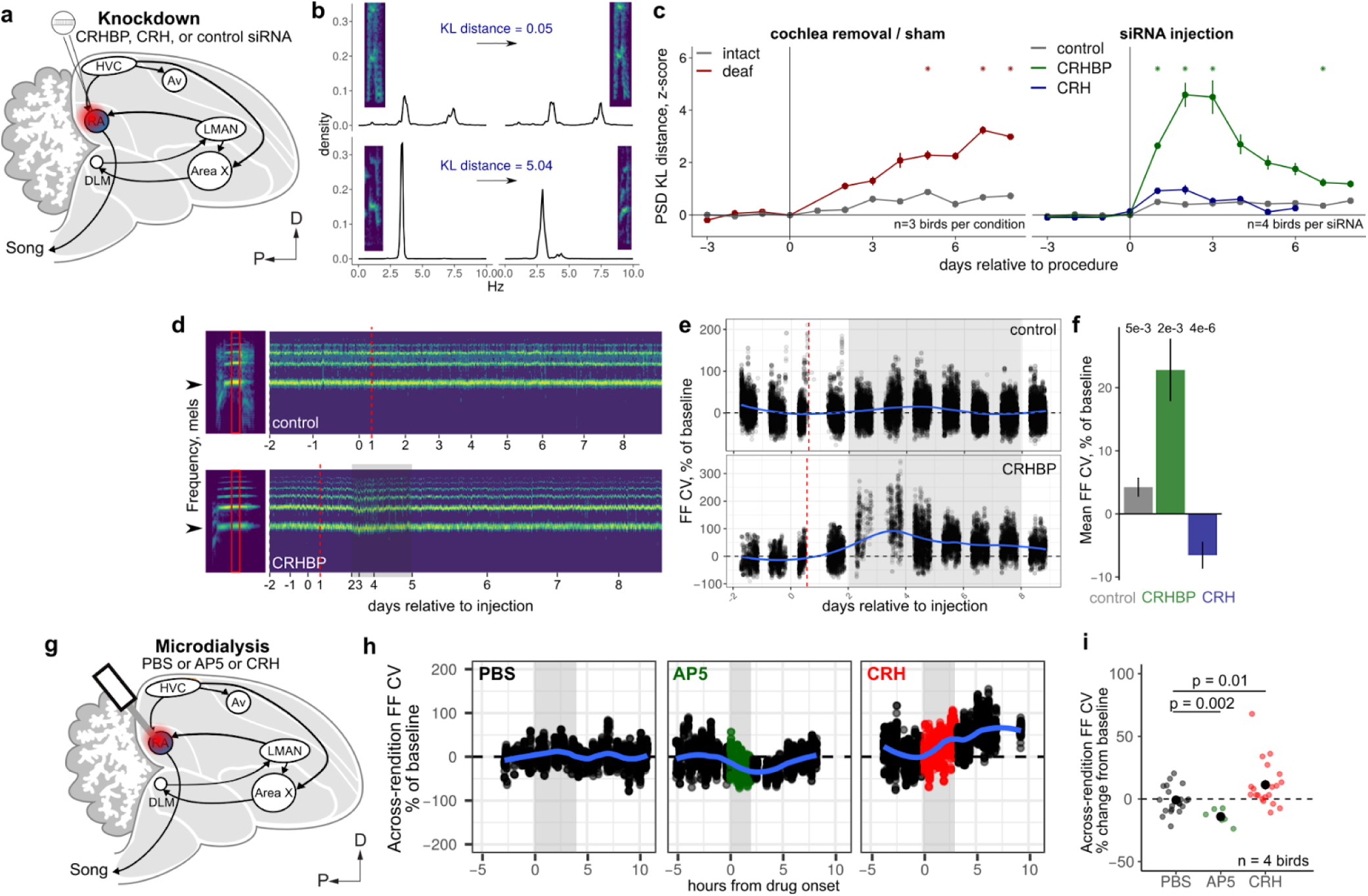
Modulation of the CRH pathway has bidirectional effects on song variability. **(a)** Transient siRNA knockdown of CRHBP and CRH in the song motor output nucleus RA. **(b)** Example calculation of KL distance between syllable power spectral densities. Top example presents a syllable that is relatively unchanged and yields a small KL distance. The syllable in the bottom example is more divergent, yielding a higher KL distance. **(c)** Power spectral density (PSD) Kiebler-Lullbeck (KL) distances over time following (*left*) cochlea removal or sham surgery or (*right*) injection of *CRHBP*, *CRH*, or control siRNAs into RA. The PSD KL distance for each syllable was calculated relative to the average PSD for the same syllable in the pre-procedure baseline period. Values were then z-scored relative to the mean and standard deviation of PSD KL distances in the baseline period. * p < 0.01, linear mixed effects model comparing the difference from baseline. **(d)** Influence of CRHBP knockdown on syllable variability. *Left,* Spectrograms of two syllables sung during the pre-injection period, one from a bird injected with a control siRNA and one from a bird injected with a siRNA targeting CRHBP. Spectrograms were averaged along the time axis within red boxed regions shown on the examples and plotted over time. Arrowheads indicate the fundamental frequency. Vertical dashed red line indicates the day after siRNA knockdown. Shaded gray box highlights a period of high syllable variability post CRHBP knockdown. Tick marks indicate the beginning of each day. Shown is 20% of the total number of syllables. **(e)** Fundamental frequency variability over time for the example syllables shown in panel (D). Each point is the FF coefficient of variation calculated across a window including the fundamental frequencies of a given syllable and the 10 syllables preceding it. Vertical dashed red line indicates the time of siRNA knockdown. Shaded gray box indicates the period used to calculate post-knockdown summary statistics in panel (F). **(f)** Influence of CRH pathway knockdowns on fundamental frequency (FF) across-rendition variability. For each bird, syllable, and period (pre vs. post-knockdown), the across-rendition coefficient of variation was computed. Values were then normalized within each bird and syllable to give a percent change relative to pre-knockdown values. These normalized values were then averaged across birds and syllables. Shown here are the normalized post-knockdown values. Pre-knockdown data includes at least 2 days of song prior to injection. Post-knockdown data includes days 2-7 following injection. Error bars are standard errors across birds. A linear mixed-effects model (see Methods) was fit to the data to obtain p-values. **(g)** Chronic reverse microdialysis of CRH and the NMDA receptor inhibitor AP5 to the song motor output nucleus RA to assess the real-time influence of CRH on song variability. **(h)** Influence of AP5 and CRH on across-rendition fundamental frequency (FF) coefficient of variation (CV). Depicted are three example days of reverse microdialysis of either PBS, AP5, or CRH. Each point is the FF CV calculated across the 5 syllables flanking each syllable on either side. Gray region indicates the period of drug delivery. For the PBS control, a syringe with fresh PBS was swapped in for a 4 hour period to mimic the syringe change during drug delivery. **(i)** Summary of FF CV changes across birds and syllables. Shown are measures for syllables with easily computed FF (harmonic stacks) that had consistent reductions in CV following AP5 treatment. Each dot represents the average FF CV within the treatment period for a given syllable, normalized to the average FF CV in the pre-treatment period (from 2 hours before drug onset to drug onset). A linear mixed-effects model (see Methods) was fit to the data to obtain Benjamini-Hochberg adjusted p-values.

To assay the effects of knockdowns on song variability, we examined how the mean and coefficient of variation (CV) for the fundamental frequency (FF) of each harmonic stack in a bird’s song changed following knockdown (Fig. 7D-F and Fig. S8F). Both *CRHBP* and *CRH* knockdown resulted in slight reductions in FF mean (*CRHBP*, -0.64 ± 0.33%; *CRH,* -0.94 ± 0.14%, Fig. S8F). In contrast, *CRHBP* and *CRH* knockdowns had opposite effects on FF variability. Knockdown of *CRHBP* increased FF CV during a 2-7 day period following knockdown (23 ± 5%) while knockdown of *CRH* reduced FF CV during this period (-6.5 ± 2.1%, Fig. 7D-F).

We next examined how elevating levels of CRH in the song system influences song variability by delivering CRH neuropeptide to RA using reverse microdialysis (n=4 birds, Fig. 7G). To validate microdialysis probe targeting, we first delivered the NMDA receptor antagonist AP5 into RA. This manipulation has previously been shown to reduce the variability of FF across syllable renditions, corresponding to the degree of variation in the mean FF from one syllable rendition to the next (Charlesworth et al., 2012; Olveczky et al., 2005; Tian and Brainard, 2017; Warren et al., 2011). As previously observed, AP5 delivery reduced the variability of FF across renditions (Fig. 7H and I, -13.5 ± 1.5% decrease with AP5 relative to baseline vs. 1.2 ± 1.7% increase with PBS relative to baseline, p = 0.002, linear mixed-effects), indicating the efficacy of our targeting of drug delivery to RA. After recovery, we then tested the effects of delivering CRH through the same microdialysis probes. Consistent with our model, we found that delivery of CRH increased the variability of FF across syllable renditions (Fig. 7H and I, 8.8 ± 2.8% increase with CRH relative to baseline vs. 1.2 ± 1.7% with PBS relative to baseline; p = 0.01, linear mixed-effects). Combined, these results indicate that the CRH pathway modulates song variability, with CRHBP acting to decrease and CRH acting to increase song variability.

## Discussion

### The ability to reliably execute learned motor skills is a critical function of the nervous system

How molecular properties of sensorimotor circuits modulate the stability of motor performance is poorly understood. Songbirds, with their complex learned vocalizations and a neural circuit that is dedicated to its performance and learning, are a powerful model through which to link candidate molecular regulators to behavioral output. Here, we examined the influence of neuropeptide signaling systems, prominent modulators of neural circuit function in other models, on birdsong. We assessed the regional and cellular expression profiles of neuropeptides, their receptors, and their modulators across the song system, the relationships between their expression and song stability and performance, and the functional influence of one candidate pathway, the CRH pathway, on song variability. Combined, our results point to a central role for neuropeptide signaling in regulating birdsong performance and suggest that neuropeptide activity in central motor circuits influences the precision of motor output.

### Role of neuropeptides in the local modulation of motor performance

Past work in a number of other systems has demonstrated a critical role for neuropeptides in regulating motor output. Neuropeptides regulate the output of central pattern generators in the feeding systems of the mollusc *Aplysia californica* as well as crabs and lobsters by modulating the influence of certain neuronal populations on circuit activity (Nusbaum and Blitz, 2012). Similarly, in *C. elegans*, neuropeptides act on motor circuits to modulate the rate and reversal frequency of locomotion, in effect regulating the balance between exploratory and exploitative feeding behavior (Bhat et al., 2021). However, the roles of neuropeptides in regulating motor performance in vertebrates are poorly studied, particularly as it relates to how these signaling systems contribute to the regulation of cortical and pallial circuits.

The specialized expression patterns of neuropeptides and their receptors in song regions relative to adjacent non-song circuits suggests that local neuropeptide signaling has a key role in regulating the song performance. Furthermore, these signaling systems are poised to have diverse effects on circuit activity due to the specificity of their expression in defined neuron populations in HVC and RA. Indeed, we find here that many of the genes whose expression is altered during song destabilization and singing are neuropeptides or neuropeptide regulators.

### Roles of CRH and CRHBP in modulating motor variability and plasticity

Our data indicate that corticotropin-releasing hormone binding protein (CRHBP) and its target the neuropeptide corticotropin-releasing hormone (CRH) influence features of song stability, including rendition-to-rendition variability. CRH (also known as CRF) and CRHBP are central factors in regulating hormonal stress responses through the hypothalamic–pituitary–adrenal axis (Ketchesin et al., 2017). However, this signaling system has long been recognized as a modulator of a variety of neurophysiological properties in neurons of the central nervous system (Gallagher et al., 2008). CRH increases the excitability of neurons in the prefrontal cortex, hippocampus, and cerebellum (Aldenhoff et al., 1983; Blank et al., 2003; Fox and Gruol, 1993; Kratzer et al., 2013; Li et al., 2016), potentiates NMDA receptor currents in dopaminergic neurons (Ungless et al., 2003), and can prime long-term potentiation in the hippocampus (Blank et al., 2002). In addition to these stimulating effects, prolonged elevated levels of CRH have also been shown to lead to dendritic spine loss (Chen et al., 2008; Y. Chen et al., 2013; Wang et al., 2013). Notably, CRH in the central amygdala has been shown to promote the acquisition of fear memory by increasing the gain of weak fear-inducing stimuli (Sanford et al., 2017), suggesting that the neuropeptide primes neural circuits for plasticity. Recent work has also demonstrated that local CRH release in the medial prefrontal cortex regulates novelty seeking in mice (Riad et al., 2022).

CRHBP is a secreted protein (Blanco et al., 2011) and modulates CRH signaling, primarily acting as a ligand sink and preventing the binding of CRH to its receptors (Kemp et al., 1998). This inhibitory role is supported by studies showing that elevated CRHBP levels reduce CRH signaling (Cortright et al., 1995; Huising et al., 2008) and that ligands that inhibit the binding of CRH to CRHBP amplify the electrophysiological and neural effects of the neuropeptide (Behan et al., 1995; Li et al., 2016). However, CRHBP function may be more complicated as there is evidence that it can promote CRHR2 signaling by acting as a cofactor for CRH binding (Ungless et al., 2003) and increasing receptor surface expression (Slater et al., 2016).

CRHBP’s inhibitory role and CRH’s role in increasing the excitability and synaptic transmission of target neurons suggests that CRHBP acts to limit the influence of CRH on neural activity in the song system. Extending from the electrophysiological effects of CRH in other forebrain neurons, we hypothesize that CRH potentiates NMDA receptor currents in the output motor nucleus RA. Past work has demonstrated that the anterior forebrain pathway (AFP) increases song variability through its projection from the AFP output nucleus LMAN to RA (Kao et al., 2005; Olveczky et al., 2005; Kao and Brainard, 2006; Aronov et al., 2008; Andalman and Fee, 2009; Stepanek and Doupe, 2010; Ölveczky et al., 2011; Warren et al., 2011). Inputs from LMAN to RA projection neurons predominantly occur through NMDA receptor-mediated transmission (Mehaffey and Doupe, 2015; Mooney and Konishi, 1991; Stark and Perkel, 1999) and blockade of these receptors reduces rendition-to-rendition song variability (Olveczky et al., 2005; Warren et al., 2011). Consistent with this model, we show that reducing CRHBP expression in RA increases song variability while reducing CRH expression in RA decreases song variability. Correspondingly, increasing levels of CRH in RA results in increased song variability.

### Dynamic regulation of CRHBP

Recent analyses of single-cell expression data from the mammalian neocortex found that neuropeptidergic networks show dense expression patterns, such that almost all neuron types express at least one neuropeptide precursor and one cognate receptor (Smith et al., 2019). These dense networks are also found in turtle and songbird neuronal populations, suggesting that this structure is a deeply conserved feature of pallial neural circuits (Smith, 2021). Consistent with this conservation, we found that the components of the CRH pathway are expressed in similar neuronal populations in songbirds, mice, and turtles. CRHBP is expressed in MGE-derived GABAergic interneurons, such as *Sst-* and *Pvalb-* class neurons, while CRH is expressed in *Vip-*class interneurons (with the exception of turtle in which CRH is expressed in a range of neurons). In contrast, CRH receptors were predominantly expressed in glutamatergic projection neurons. This pattern — neuropeptides in GABAergic neurons and receptors in glutamatergic neurons — is shared across a number of neuropeptidergic systems (Smith, 2021), forming a conserved motif across species.

Interestingly, the modulatory logic of the CRH system is consistent with the roles of each neuron type in local microcircuitry. In mammalian neocortex, *Vip-*class interneurons inhibit other interneurons resulting in the disinhibition of glutamatergic neurons; these neurons also express *CRH*, which increases the neural activity of glutamatergic neurons. Likewise, *Sst/Pvalb-*class interneurons inhibit glutamatergic neurons, and *Sst/Pvalb* neurons also express *CRHBP,* which limits the stimulating effects of *CRH*. These similarities are consistent with a model in which both fast synaptic and slow modulatory signaling act in parallel to concordantly regulate pallial neural circuits on different time scales.

CRH/CRHBP signaling is likely one among several neuropeptide systems that in concert influence song performance. Our transcriptional analysis of the song system and surrounding brain regions indicates that other signaling pathways are enriched in birdsong-dedicated regions and may be modulated in conjunction with song learning and plasticity. Like the CRH pathway, these pathways may play roles in establishing neuromodulatory states that facilitate or limit song stability. Targeting these pathways using genetic and pharmacological strategies, combined with song analysis and neurophysiological characterization, would serve as a tractable model for understanding their roles in modulating sensorimotor skill performance.

## Materials and Methods

### Animal care and use

All Bengalese finches were from our breeding colonies at UCSF or were purchased from approved vendors. Experiments were conducted in accordance with NIH and UCSF policies governing animal use and welfare.

### Definition of neuropeptide-associated genes

An initial list of neuropeptide-associated genes was generated by compiling the genes present in the following gene ontology terms: GO:0005184, neuropeptide hormone activity; GO:0071855, neuropeptide receptor binding; GO:0160041, neuropeptide activity; GO:0008188, neuropeptide receptor activity; GO:0001635, calcitonin gene-related peptide receptor activity; GO:0015056, corticotrophin-releasing factor receptor activity; GO:0004995, tachykinin receptor activity; GO:0038170, somatostatin signaling pathway. To this list, the following neuropeptides, receptors, and neuropeptide regulators were added: CRHBP, NPY2R, NPY5R, OPRK1, OPRM1, PDYN, TAC1, TAC3, CCKBR, VIPR1, VIPR2, NTSR1, ADCYAP1R1. In total this yielded 80 genes.

### Differential expression analysis

Gene-sample count matrices were filtered to remove lowly expressed genes, defined as having a total number of reads across samples less than the number of samples divided by eight (the number of brain regions assayed). For each sample we also calculated the ‘cellular detection rate’ (CDR, the number of genes detected in a given sample), previously shown to substantially influence differential expression analysis on single-cell RNA-sequencing samples (Finak et al., 2015). Low quality samples were defined as having a CDR less than 30% of the total number of genes in the reference annotation (18,674 genes). Normalization factors were calculated using the function *calcNormFactors* from the R package *edgeR* v3.31.4 and the “TMMwsp” method. The count matrix, these normalization factors, and a design matrix were then provided to the function *voom* from the *limma* package v.3.48.3 (Law et al., 2014; Ritchie et al., 2015). The design matrix was specified as:

> ∼ 0 + position + cdr_scaled + frac_mito_scalewhere ‘position’ is an indicator for brain region, ‘cdr_scale’ is CDR, and ‘frac_mito_scale’ is the fraction of reads mapping to mitochondrial genes in a given sample.

For regressions against singing-related variables the design matrix was specified as:

> ∼ 0 + position + position:num_songs_on_euth_date_cut3 + cdr_scaled + frac_mito_scaleor

> ∼ 0 + position + position:nsongs_last_two_days_cut3 + cdr_scaled + frac_mito_scalefor the ‘2 hour’ and ‘2 day’ analysis of singing amount. Song-related variables were cut into three groups (‘cut3’) to generate low, mid, and high categories. Variables with ‘scale’ in their names were mean-subtracted and standard deviation-normalized. To calculate the within-bird correlation between samples, the resulting voom object was passed to *duplicateCorrelation* with block specified as the bird ID. To fit the above model, the voom object, design matrix, and the consensus within-bird correlation were input to function *lmFit* from limma. Coefficient estimates and standard errors for each coefficient were calculated using function *contrasts.fit*, the function *eBayes* was used to compute moderated t-statistics and p-values, and the function *topTable* was used to adjust p-values using the Benjamini-Hochberg method. Genes were considered differentially expressed if their adjusted p-values were less than 0.1.

Expression estimates and standard errors for a given bird and brain region were computed using a regression approach with a design matrix specified as:

> ∼0 + position:tags + cdr_scale + frac_mito_scalewhere ‘position’ is an indicator for brain region, ‘tags’ is the bird ID, ‘index2’ is a categorical variable indicating the sequencing run, and ‘cdr_scale’ and ‘frac_mito_scale’ are as described above. Standard errors were extracted from the linear fit model ‘fit’ as: sqrt(fit$s2.post) * fit$stdev.unscaled.

### Gene set enrichment analysis

Gene Ontology lists were obtained from the Molecular Signatures Database (set C5, version 7). Gene set enrichment analysis was performed using the R package *fgsea* v1.18.0 (Korotkevich et al., 2021). T-statistics from *voom* regression or gene module membership scores from *MEGENA* (see (Colquitt et al., 2023)) were input into the function *fgseaMultilevel* (minSize = 20, maxSize = 200). Resulting pathways were filtered for those with an adjusted p-value less than 0.2 and similar pathways were pruned using *collapsedPathways* (pval.threshold = 0.01 or 0.05).

### Neuropeptide-receptor analysis

Neuropeptide ligand and receptor interactions were analyzed using CellChat, v2.1.2 (Jin et al., 2021, 2023). The communication probability between two cell types through a particular ligand-receptor pair is a function of the known interactions among ligands and receptors, the expression level of these pairs across cell types, and the relative abundance of each cell type. This measure was calculated on the sc/snRNA-seq dataset from (Colquitt et al., 2021) split by brain region (HVC and RA) using the function *computeCommunProb* (type = “truncatedMean”, trim = 0.05, population.size = T). Aggregated cell-cell communication plots were generated by summing the contributions of each neuropeptide pathway within a given cell-cell comparison (e.g. the communication probabilities for all neuropeptide pathways between RA-Glut-1 and Astro-1 were summed to yield a single value). Similarly, the relative communication probability of a particular ligand-receptor pair relative to other pairs was computed by summing its communication probabilities across all cell-cell comparisons then normalizing by the sum of values across all ligand-receptor pairs.

### Fluorescent in situ hybridization (FISH)

FISH was performed using the hairpin chain reaction system from Molecular Instruments. Birds were euthanized using isoflurane, decapitated, and debrained. Brains were flash-frozen in -70 °C dry ice-chilled isopentane for 12 seconds within 4 minutes from decapitation then stored at -80 °C. Fresh-frozen brains were cryosectioned at 16 μm onto SuperFrost slides (Fisherbrand) chilled in the cryochamber then melted onto the slide using a warmed metal dowel. Slides were then transferred to -80 °C for storage. For the FISH, slides were transferred from -80 °C to slide mailers containing cold 4% PFA and incubated for 15 minutes on ice. Slides were washed three times for 5 minutes using DEPC-treated PBS + 0.1% Tween-20, dehydrated in 50%, 70%, and two rounds of 100% ethanol for 3-5 minutes each round, then air dried. Slides were then transferred to a SlideMoat (Boekel Scientific) at 37 °C. 100 μL of v3 Hybridization Buffer (Molecular Instruments) was added to each slide, which were then coverslipped and incubated for 10 minutes at 37 °C. Meanwhile, 2 nM of each probe was added to 100 μL Hybridization Buffer and denatured at 37 °C. Pre-hybridization buffer was removed, 100 μL of probe/buffer was added, slides were coverslipped and incubated overnight at 37 °C. The next day, coverslips were floated off in Probe Wash Buffer (PWB, 50% formamide, 5x SSC, 9 mM citric acid pH 6.0, 0.1% Tween-20, 50 μg/ml heparin), then washed in 75% PWB/25% SSCT (5x SSC, 0.1% Tween-20), 50% PWB/50% SSCT, 25% PWB/75% SSCT, 100% SSCT for 15 minutes each at 37 °C. This was followed by 5 minutes at room temperature in SSCT. Slides were incubated in 200 μL of Amplification Buffer (provided by company) for 30 minutes at room temperature. Alexa fluor-conjugated DNA hairpins were denatured for 90 seconds at 95 °C then allowed to cool for at least 30 minutes in the dark at room temperature. Hairpins were added to 100 μL amplification buffer, applied to slides, and incubated overnight at room temperature. The following day, slides were washed in SSCT containing 1 ng/mL DAPI for 30 minutes at room temperature, then SSCT for 30 minutes at room temperature, followed by a final 5 minutes in SSCT at room temperature. Prolong Glass Antifade Medium (Thermofisher) was added to each slide then coverslipped. Sections were imaged on a confocal microscope (Zeiss 710) using a 20X objective.

### FISH quantification

Image quantification was performed using CellProfiler v4.0.4 (Stirling et al., 2021). DAPI-stained nuclei were first identified using the ClassifyPixels-Unet module. Areas corresponding to cells were estimated by extending nuclei boundaries by 5 pixels. Then signal puncta for each channel were identified and their intensities were measured. For each cell and each channel, we calculated the summed signal intensity of overlapping puncta divided by the cell area. To test for significant differences in gene expression between hearing and deaf birds, a linear mixed effects model was fit using function *lmer* from R package *lmerTest v3.1-3* (Kuznetsova et al., 2017) for each target gene and brain region as ‘intensity ∼ condition + (1|bird)’ where ‘condition’ is developmental stage, hearing, or deaf and ‘(1|bird)’ is the per-bird grouping factor. P-values were obtained by comparing this model with a reduced model ‘intensity ∼ (1|bird)’ using ANOVA.

### mRNA transcript preparation and quantification

Total RNA was purified according to the TRIzol protocol, resuspended in water, and stored at -80C. 1-10 ng of total RNA was treated with 1μL ezDNase (ThermoFisher) in 10 μL reactions for 2 minutes at 37C then placed on ice. Reverse transcription was performed using the SuperScript IV VILO Master Mix (ThermoFisher), adding 4 ul of Master Mix to the DNase-treated total RNA and 6 ul water. Samples were then incubated at 25C for 10 minutes, 50C for 10 minutes, then 85C for 5 minutes. cDNA was stored at -20C. To quantify relative transcript abundance, we used quantitative PCR via the TaqMan system (ThermoFisher). TaqMan probes were designed against Bengalese finch *CRHBP*, *CRH, SST*, and *PPIA* (housekeeping gene control). CRHBP, CRH, and SST probes were designed with FAM dye label and PPIA with VIC label. For each sample, either CRHBP, CRH, or SST was combined with PPIA to serve as a loading control. Assays were performed on a QuantStudio 6 Real-Time PCR machine (Applied Biosystems).

### In vitro knockdown

Custom 21 nucleotide siRNAs (Dharmacon) were designed against CRHBP and CRH transcript sequences (Bengalese finch genome annotation, lonStrDom1, RefSeq assembly GCF_002197715.1) using the Dharmacon design tool. A control siRNA was also designed that had similar GC content as the CRHBP siRNA and targeted no known Bengalese finch transcript.

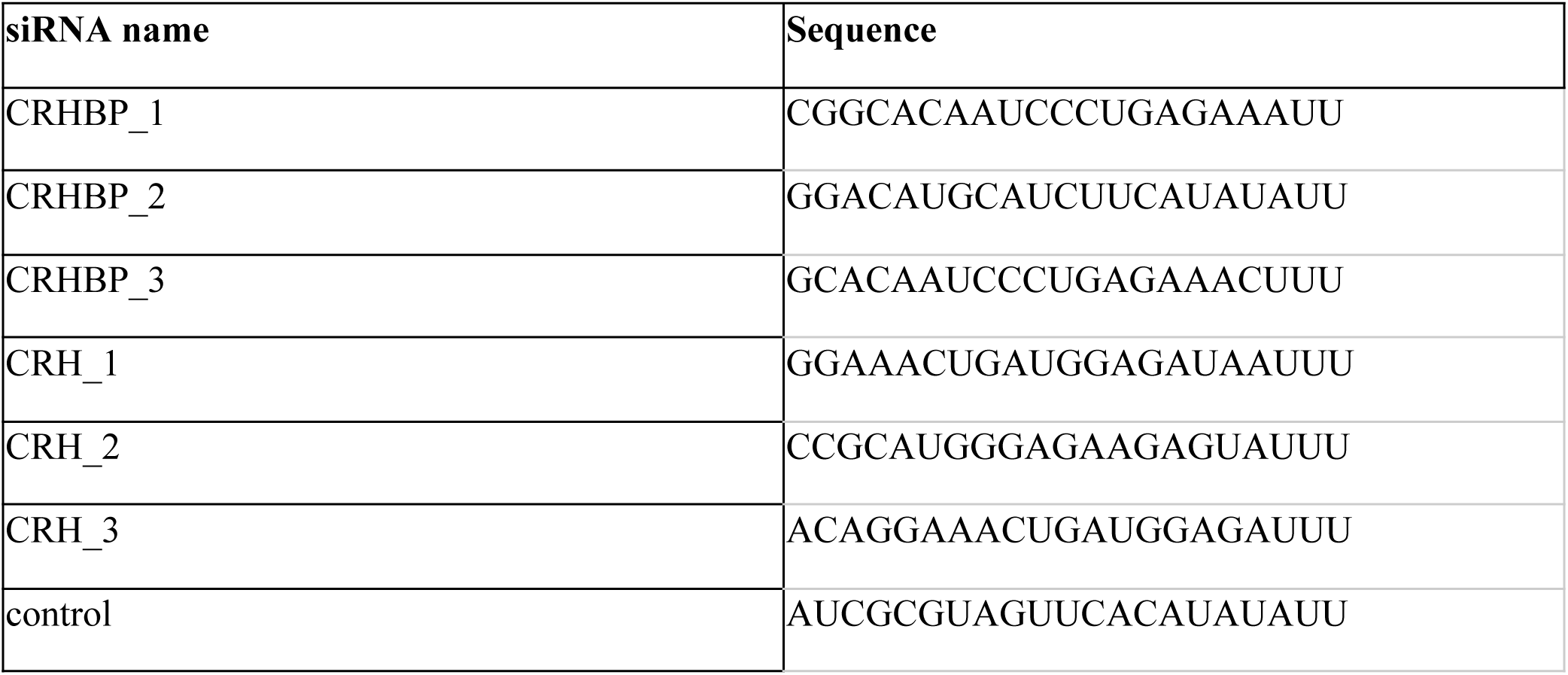

To validate knockdown efficacy in vitro, we transfected siRNAs into Bengalese finch primary neural culture. Before the preparation, Neurobasal A complete (NBAC) was prepared as 95 mL Neurobasal A (ThermoFisher), 1 mL 100x N-2 Supplement (Thermofisher), 2 mL 50x B-27 Supplement (ThermoFisher), 1 mL GlutaMAX Supplement (Thermofisher), and 1 mL 10,000 U/mL Penicillin-Streptomycin (ThermoFisher). NBAC was filtered through a 0.22 μm syringe filter and stored at 4 °C for no more than 15 days. Juvenile birds (1-14 days post hatch) were decapitated, and brains were dissected into cold Hank’s Balanced Salt Solution (HBSS, no phenol red, with magnesium and calcium, ThermoFisher) in a dish on ice. Tissue was then transferred to a chilled eye glass dish containing 500 μL HBSS and minced using forceps. The minced tissue was then transferred to a 1.5 mL tube and kept on ice. Papain solution was prepared by adding a small amount of lyophilized papain (Worthington) to 1 mL of HBSS. Tissue was dissociated by replacing the HBSS media in the tube containing the brian tissue with papain solution (20 U/mL in HBSS), followed by incubation at 37 °C for 45 minutes. Tissue was then tritruated 20 to 30 times using a 5 mL serotological pipette and centrifuged at 1200 RPM for 5 minutes at room temperature. Supernatant was removed and replaced by 1 mL of 7 mg/mL ovomucoid trypsin inhibitor in HBSS (Worthington). Samples were then centrifuged at 1200 RPM for 5 minutes at room temperature, supernatant was removed, and 8 mL of NBAC warmed to 37 °C was added. Samples were then triturated 40 times using fire-polished glass pasteur pipettes. Debris was allowed to settle then the supernatant was filtered through a 100 μm cell strainer. Cell suspensions were counted, diluted to 0.5 millions cells / mL, then distributed across 24-well plates with 1 mL per well. Cells were incubated at 37 °C in 5% CO_2_ for 4 hours to allow plate attachment, then 50% of the media in each well was replaced with fresh media. 50% of the media was again replaced with fresh media 3 days later.

Once neural processes formed (after ∼1 week) the cell cultures were used to test siRNA molecules. For each siRNA tested, knockdown experiments were conducted in duplicate using the RNAiMAX (Life Technologies) transformation reagent. For each siRNA, 25 μL of Opti-MEM Medium (Life Technologies) was mixed with 1.5 μL of RNAiMAX reagent. Separately, 25 μL of Opti-MEM Medium was mixed with 5 pmol annealed siRNA molecules (Dharmacon). These two mixtures were then combined and incubated at room temperature for 5 minutes. Following incubation, 10 μL of the RNAiMAX/siRNA mixture was added to each of two wells of neural cell culture. Cultures were then incubated at 37°C and 5% CO_2_ for 2 days. After incubation, RNA from each well was purified, converted to cDNA, and quantified using the protocol described in mRNA transcript preparation and quantification (above). The mRNA expression level of the ‘target’ gene was compared between cells exposed to experimental siRNA molecules and cells exposed to a control siRNA molecule.

### In vivo knockdown

To knockdown CRHBP and CRH in RA, siRNA stereotactically delivered using a cationic transfection reagent, BrainFectIN (OZ Biosciences). Adult male birds were anesthetized using isoflurane then placed in stereotax with a beak angle of 32 degrees. A midline incision was made through the skin of the skull to expose Y0 and the surface position of RA was marked at 0.5 mm posterior to Y0 and 2.3 mm lateral from Y0 on both hemispheres. Craniotomies were opened at these regions, and the dorsal ventral position of RA was determined using a carbon fiber electrode to identify regions with high tonic activity that is characteristic of RA. siRNA complex was then prepared at room temperature by mixing 1 μL of 2 μg/μL siRNA with 3 μL of BrainFectIN followed by mixing 20 times by pipette. For CRHBP knockdown, CRHBP_1, CRHBP_2, and CRHBP_3 siRNAs were combined at equal concentrations.

For CRH knockdown, CRH_1 and CRH_2 were combined at equal concentrations. siRNA/BrainFectIN solution was allowed to incubate for 15-30 minutes at room temperature, then 1% FastGreen FCF was added to a final concentration of 0.1%. The solution was then loaded into a glass capillary with a 20-30 μm tip pore using a NanoJect II (Drummond). 100 nL of siRNA/BrainFectIN was injected using 43 2.3 nL pulses with 10 second pauses between pulses, followed by a 10 minute post-injection pause. This procedure was repeated for the other hemisphere using the same siRNA/BrainFectIN complex. The final injection occurred within 90 minutes of siRNA complexing. After both injections, the midline incision was sealed using VetBond (3M), and birds were allowed to recover in a sound isolation chamber.

For validation, birds were injected as above with control siRNA into one hemisphere and either CRH or CRHBP siRNAs into the other hemisphere. Birds were euthanized two days after euthanasia, brains were disssected out into ice-cold DPBS, and sliced on a vibratome at 260 μm thickness. The injected area (visible as a green dot due to FastGreen signal) was microdissected out into 1 mL TRIzol (ThermoFisher) on ice, then stored at -80 C. RNA was then purified, converted to cDNA, and quantified using the protocol described in mRNA transcript preparation and quantification (above).

### Reverse microdialysis

Guide cannulae for CMA7 or CMA8 probes were implanted bilaterally above and slightly lateral to RA (beak angle 32 degrees, ML 2.35 mm, AP 0.51 mm anterior to Y0). Craniotomies were opened at these regions, and the dorsal ventral position of RA was determined using a carbon fiber electrode to identify regions with high tonic activity that is characteristic of RA. Cannulae were then implanted 1.2 mm above the dorsal surface of RA. After one week of recovery, dialysis probes were inserted into the cannulae. Either CMA7 (molecular weight cutoffs (MWCO) of 6 kDa) or CMA8 (MWCO 20 kDa) probes. Results were consistent for both probe types, and data from experiments using each was pooled. A syringe pump was used to continuously pump PBS through the probes at a rate of 0.5 μl/min. After the birds acclimated to the setup (2-3 days following probe insertion), either 4 mM AP5 (Sigma, A8054) or 5-10 uM CRH (hrCRF, BACHEM H-2435) was delivered for 2 to 4 hours, followed by a washout with PBS.

### Song recording and preprocessing

Birds were individually housed in wire cages in sound isolation chambers. Song was recorded using Countryman Isomax microphones taped to the top of the wire cage. Microphones were connected to USB preamplifiers that were connected to a Linux workstation. Audio was recorded at a frame rate of 44,100 samples/second using a custom python script, and, to select for periods of singing, blocks of continuous sound with amplitudes above a manually set threshold were saved as 24-bit WAV files.

### Song autolabeling

The analysis of specific spectral features (e.g. fundamental frequency) was performed on syllables that were labeled using a supervised machine learning approach, called *hybrid-vocal-classifier* or *hvc* (Nicholson, 2021). For each bird, 20-50 songs were manually labeled using the Matlab software *evsonganaly*. Using *hvc*, a set of spectrotemporal features was computed for each syllable (e.g. duration, mean frequency, pitch goodness, and mean spectral flatness as defined in (Tachibana et al., 2014)). These features and the manually defined labels were provided to *hvc* to train a support vector machine (SVM) with radial basis function, with a grid search across parameters C and gamma to identify parameters with the highest classification accuracy. A set of models were then trained using these selected parameters and a random sample of training syllables, and model accuracy was tested on a held-out set of syllables. For each bird, the number of input syllables and parameters were adjusted until label accuracy reached 95-100%. The model with the highest accuracy was then used to predict labels on unlabeled songs. To select confidently labeled syllables, a prediction confidence score was calculated for each syllable as the entropy (sklearn.stats.entropy) of the classification probabilities resulting from SVM model prediction. Syllables with a prediction confidence score greater than 0.5 were retained.

### Song dimensionality reduction

To project syllable spectrotemporal structure into a reduced dimension space, we used an approach developed by (Sainburg et al., 2020) with code and example scripts obtained from https://github.com/timsainb/AVGN. Songs were first isolated from audio recordings then segmented into syllables based on amplitude threshold crossings. Spectrograms were computed for each syllable using short-time Fourier transforms (0.5 ms step size, 6 ms window size, 44,100 frames per second) and frequencies between 500 Hz and 15,000 Hz were retained. Spectrograms were converted to mel scale using a mel filter with 128 channels. Syllables were compressed in the time dimension to a framerate of 640 frames per second then zero-padded to yield a standardized dimension of 128 for a maximum syllable length of 200 ms. Before dimensionality reduction, these 128x128 spectrograms were further reduced to 16x16 matrices then flattened yielding a 256 length feature vector for each syllable. Syllable x feature vector matrices were then processed using the single-cell analysis framework *Seurat v3* (Stuart et al., 2019). Principal component analysis was performed, then Uniform Manifold Approximation and Projection (UMAP) was performed on the first 10 principal components to produce a two-dimensional reduction.

### UMAP density differences

To calculate global differences in syllable spectral structure before and after a manipulation (e.g. deafening), we split each bird’s song UMAP by day relative to the manipulation and computed two-dimensional kernel density estimates (R package *MASS* v7.3 function *kde2d*, 200 x 200 grid) for each of the these per-day plots. A baseline UMAP structure was calculated as the mean density across the 2-4 days of singing before the manipulation, then density differences were calculated by subtracting this baseline density from each per-day density plot. Positive values from each difference plot were summed to give a single statistic for each day. Significance between hearing and deaf conditions for each day was determined using a two-sided t-test.

### Fundamental frequency statistics

To calculate fundamental frequency (FF) for a given harmonic stack, we first computed the average spectrogram for 20 randomly selected syllables. We then identified a time within the syllable (relative to syllable onset) with stable FF and defined minimum and maximum frequency bounds to define a frequency band containing the FF. A short-time Fourier transform (STFT) was then calculated at this time point using function *spec* from R package *seewave* v2.1.8 (Sueur et al., 2008) (1024 window size, 44,100 frames per second). FF was estimated by interpolating the frequency spectrum on an output vector spanning the minimum and maximum frequency bounds with a resolution of 1 Hz (function *aspline* from R package *akima* v0.6-2.2). The maximum value of this interpolated frequency spectrum was taken as the FF. Rolling variability of FF was calculated as the coefficient of variation (CV, standard deviation / mean) over a set of FF values for a given syllable and the 10 prior syllables (of the same type). To compare variability relative to a baseline period, FF CV values were transformed to a percentage relative to the average variability before manipulation. Group estimates and significance values were obtained from mixed effects linear models using R package *lme4* v1.1-27.1 (Bates et al., 2015) and function *lmer* (maximum likelihood criterion). The time period (before or after manipulation) was treated as a fixed effects and bird ID and syllable were treated as random effects with syllable nested under bird ID [model in *lme4* notation: period + (1 | bird / syllable)]. P-values for fixed effect were obtained using ANOVA (Type II, Wald chi-square test statistic, R package *car* v3.0-11 function *Anova* (Fox and Weisberg, 2019)), followed by adjustments for multiple testing using Benjamini-Hochberg correction.

## Code availability

Analysis for the SLCR-seq, cellular transcriptomics, *in situ* hybridizations, and song features can be found at https://github.com/colquittlab/birdsong-neuropep.

## Data availability

SLCR-seq mapped sequencing reads, gene-by-sample count matrices, and metadata can be found at NCBI Gene Expression Omnibus (GSE200663). Single-cell RNA-seq data are available in GEO for Bengalese finches (GSE150486) and zebra finches (GSE153665).

