## Supplemental information for "An interneuronal CRH and CRHBP circuit stabilizes birdsong performance"

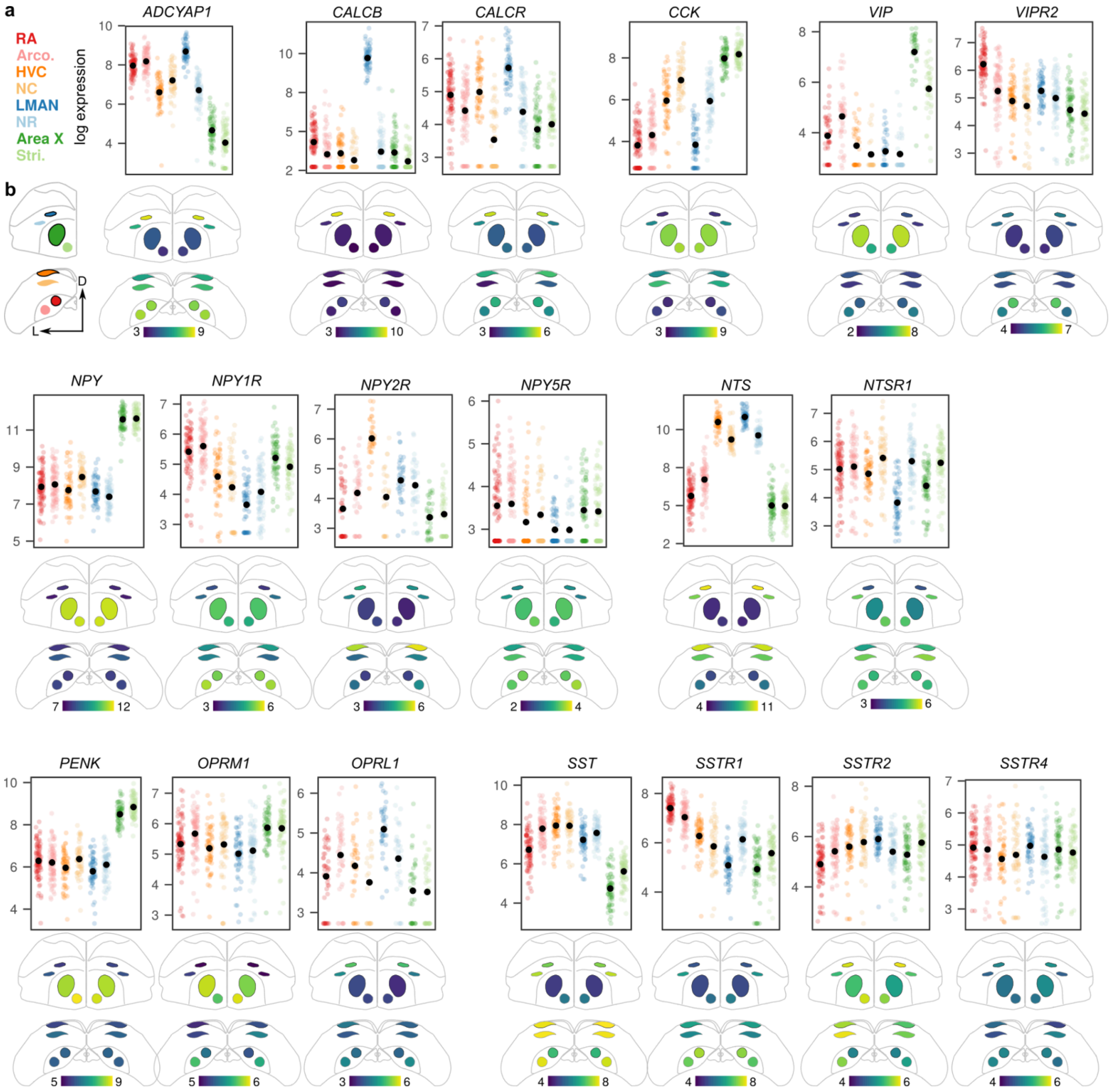


**Supplemental Figure 1. Expression of neuropeptide-associated genes across the song system.**
**(a)** Normalized log gene expression data of neuropeptide-associated genes. Plots are organized by ligand-receptor sets. Each point indicates gene expression in a single SLCR-seq sample. **(b)** Coronal anatomical atlas representation of the expression of genes plotted in (A). Each region is colored according to log gene expression value. D, dorsal; L, lateral.

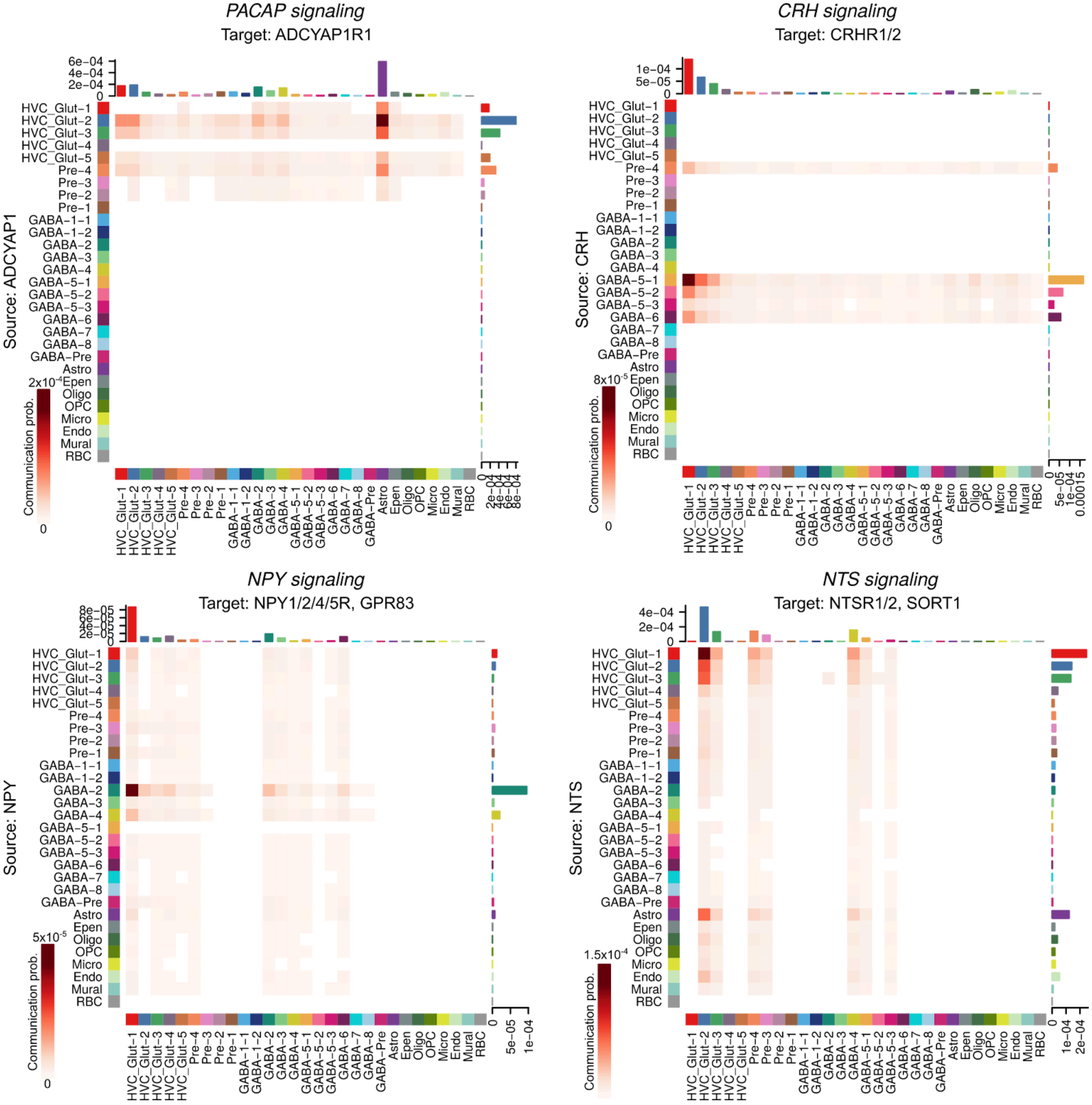


**Supplemental Figure 2. Neuropeptide communication probabilities for cell types in HVC (ADCYAP1, CRH, NPY, NTS)**

Heatmaps displaying the cell-cell communication probabilities for expressed neuropeptide pathways in HVC. Rows indicate source (ligand) cell types, and columns indicate target (receptor) cell types.

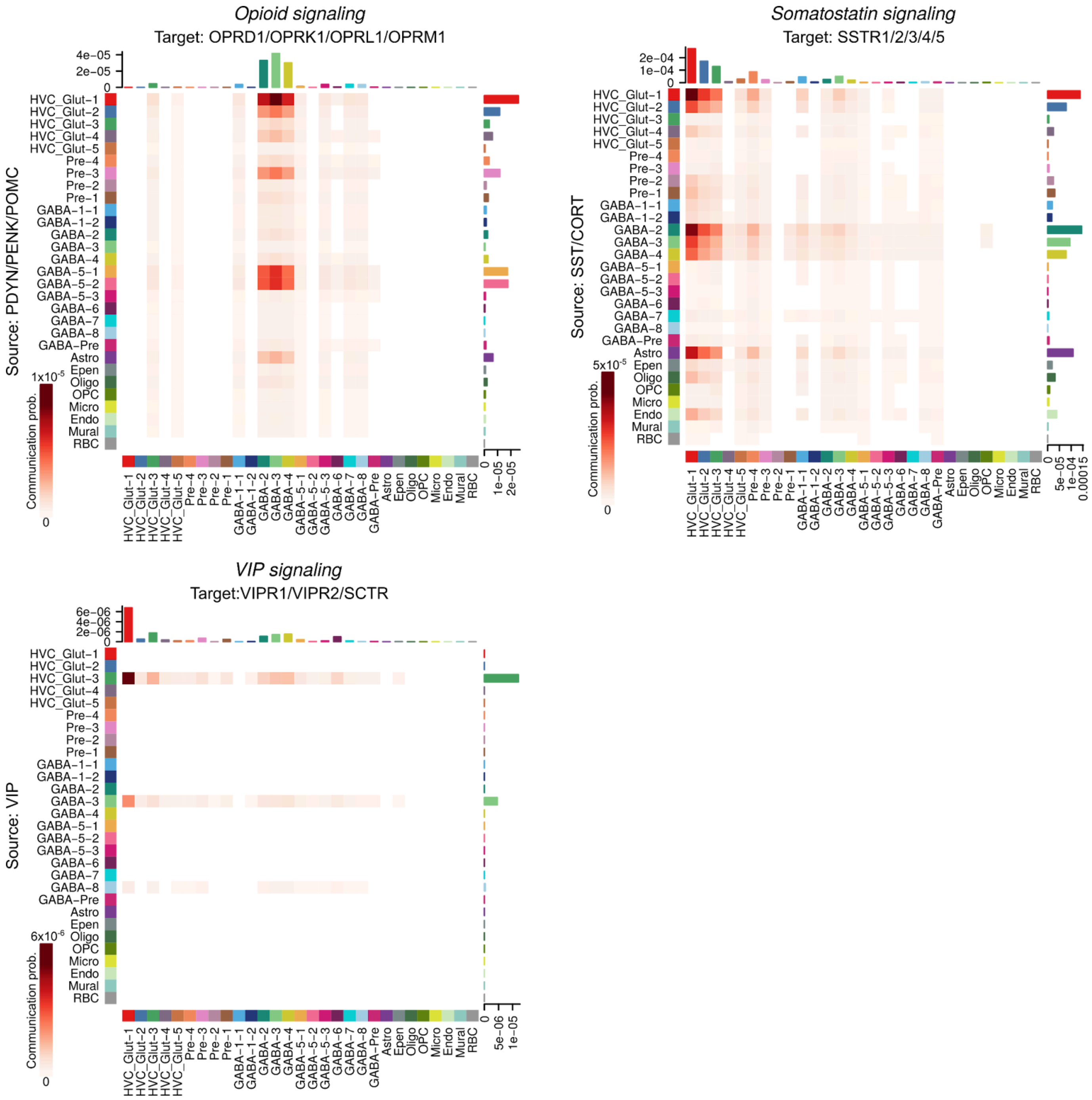


**Supplemental Figure 3. Neuropeptide communication probabilities for cell types in HVC (OPIOID, SST, VIP)**

Heatmaps displaying the cell-cell communication probabilities for expressed neuropeptide pathways in HVC. Rows indicate source (ligand) cell types, and columns indicate target (receptor) cell types.

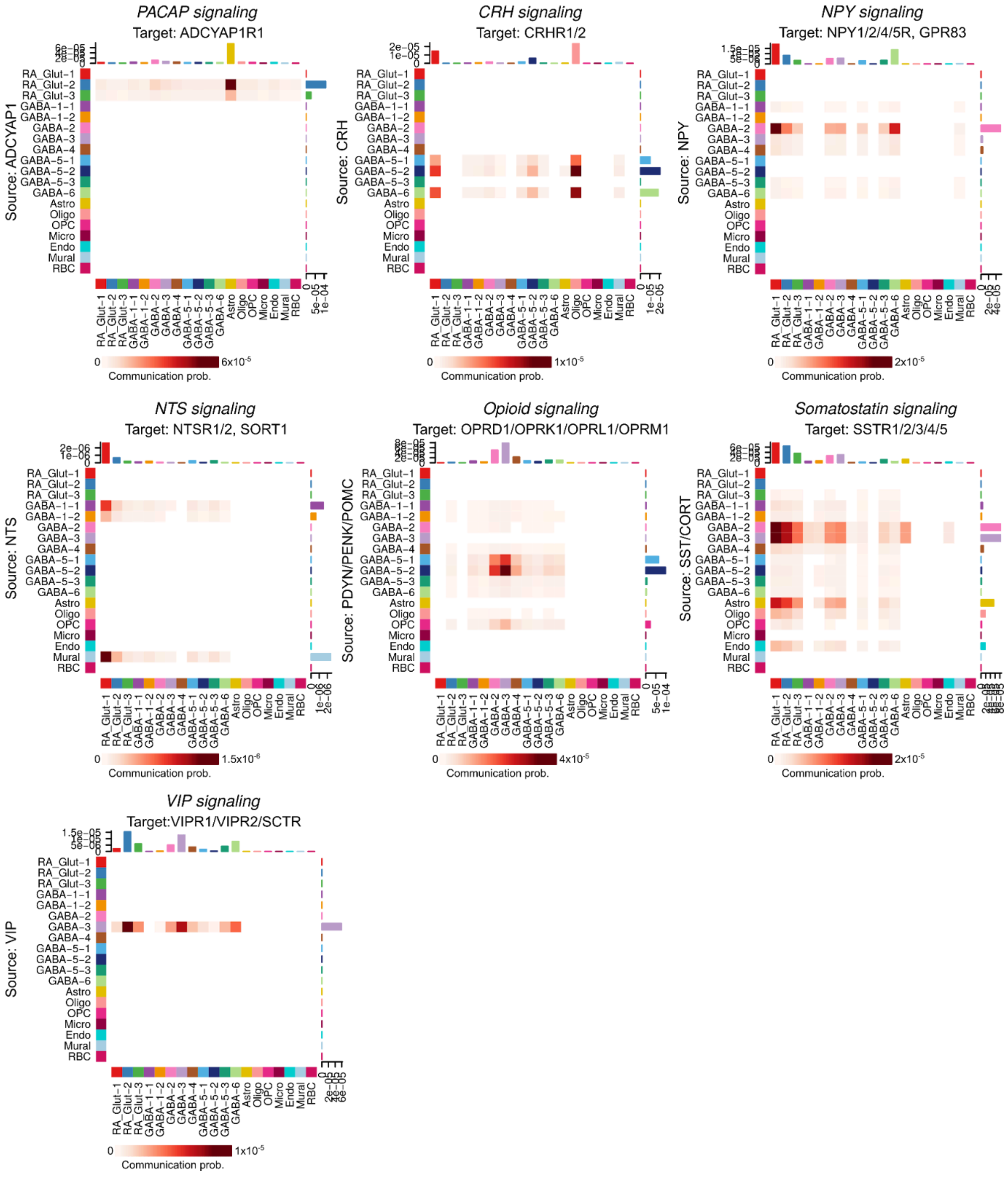


**Supplemental Figure 4. Neuropeptide communication probabilities for cell types in RA**
Heatmaps displaying the cell-cell communication probabilities for expressed neuropeptide pathways in RA. Rows indicate source (ligand) cell types, and columns indicate target (receptor) cell types.

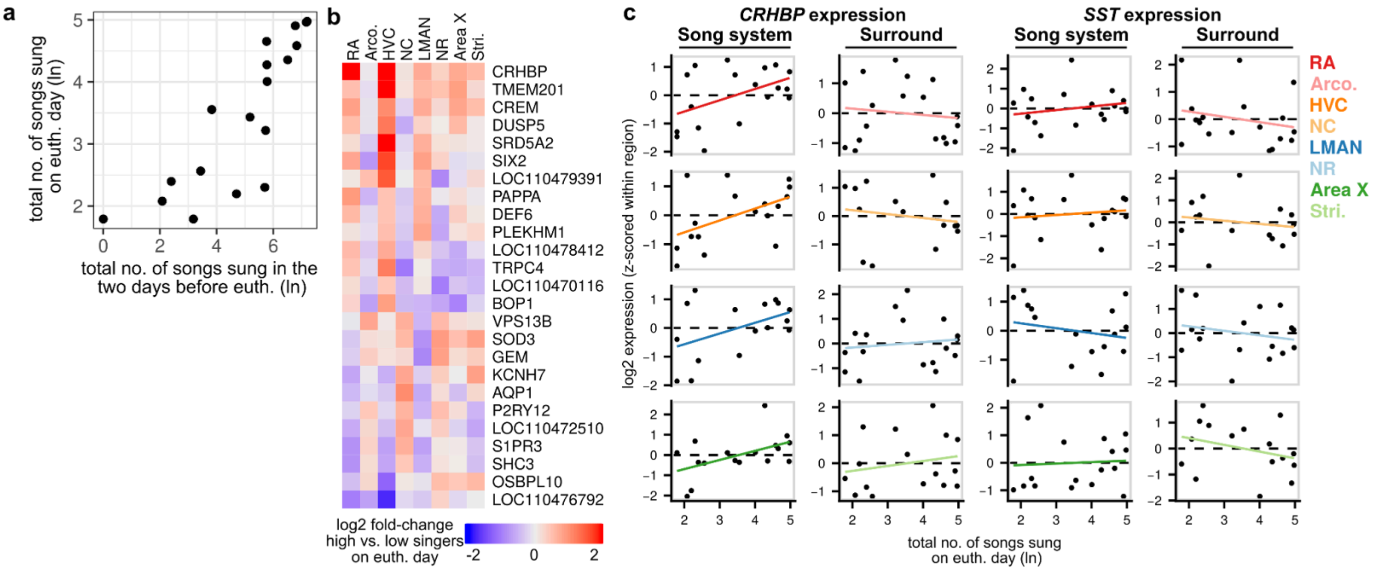


**Supplemental Figure 5. Singing-dependent gene expression in the song system**

**(a)** Relationship between the two measures of singing: total number of songs sung during the two hours before euthanasia and total number of songs sung during the two days before euthanasia. **(b)** Heatmap of genes showing significantly increased expression (adjusted p<0.1) across pallial song nuclei (RA, HVC, and LMAN) between high and low singers on the day of euthanasia. **(c)** Singing-modulated expression of *CRHBP* and *SST* across the song system and surrounding regions for the day-of-euthanasia measure.

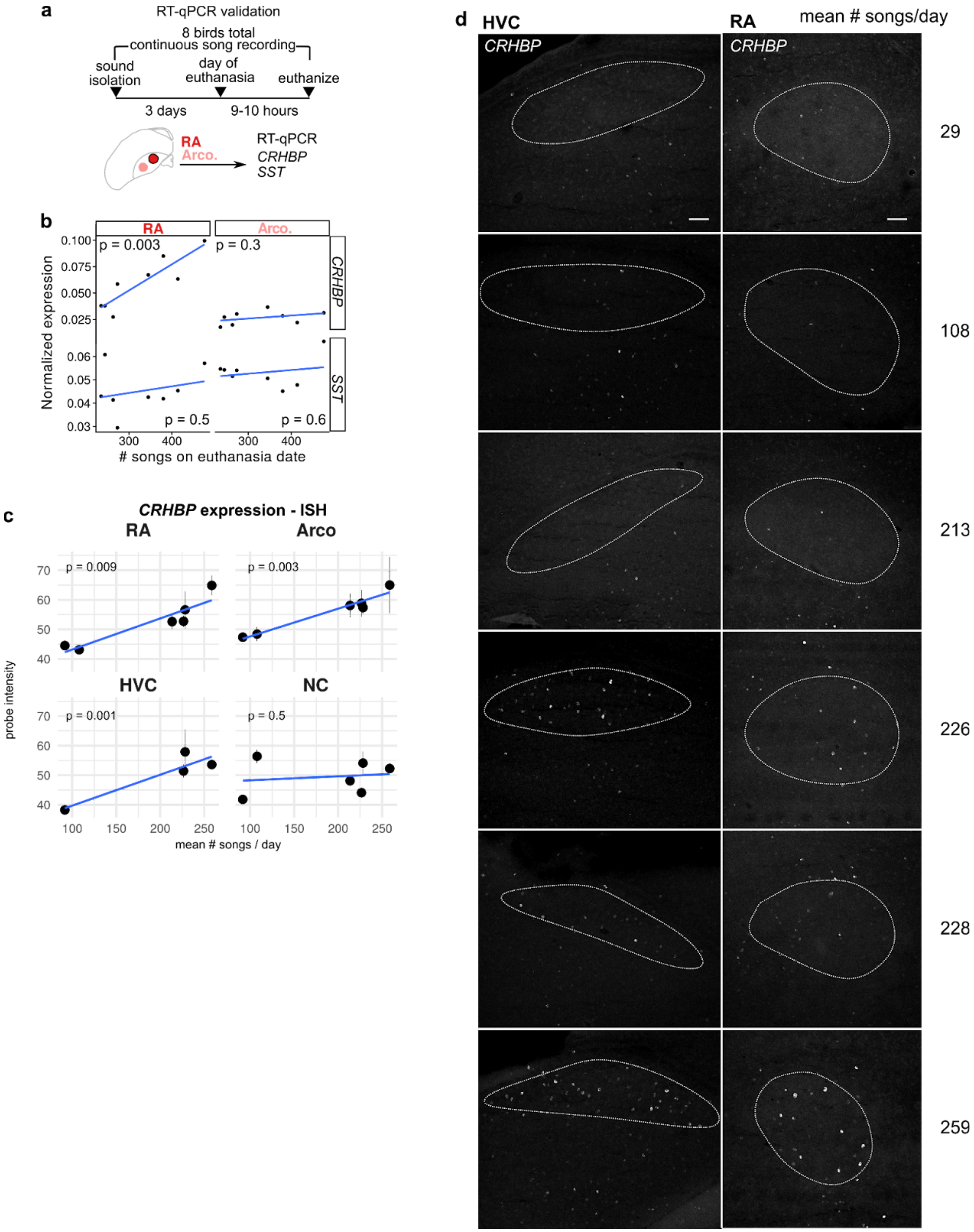

a
RT-qPCR validation
8 birds total
continuous song recording
sound isolation
day of euthanasia
euthanize
3 days
9-10 hours
RA
Arco.
RT-qPCR
CRHBP
SST
b
RA
Arco.
Normalized expression
p = 0.003
p = 0.3
CRHBP
SST
p = 0.5
p = 0.6
### songs on euthanasia date
c
CRHBP expression - ISH
RA
Arco
p = 0.009
p = 0.003
HVC
NC
p = 0.001
p = 0.5
probe intensity
mean # songs / day
d
HVC
RA
mean # songs/day
CRHBP
CRHBP
29
108
213
226
228
259

**Supplemental Figure 6. Validation of relationship between CRHBP expression in the song motor pathway and singing**

**(a)** Schematic of qPCR validation experiment. **(b)** RT-qPCR to examine the dependence of *CRHBP* and *SST* expression on the amount of singing. RA and arcopallium (Arco.) were dissected from eight birds and prepared for RT-qPCR. Across sample *CRHBP* and *SST* expression was normalized using expression of the housekeeping gene *PPIA.* P-values were calculated using linear regression between normalized *CRHBP* or *SST* expression and the number of songs sung on the day of euthanasia. **(c)** Quantification of *CRHBP* intensity in HVC and RA versus mean number of songs per day. *CRHBP*-positive cells were identified (see Methods) and the mean signal intensity calculated for each positive cell. Shown is mean±SEM probe intensity for each bird calculated from 1-2 images per nucleus per bird. Significance was calculated by performing an ANOVA comparing two mixed-effects models, one including the mean number of songs per day as a regressor and one including just the intercept term. For both models, bird identification was included as a grouping variable. **(d)** Representative *in situ* hybridizations for *CRHBP* expression in HVC and RA. Mean number of songs per day for each bird (row) is shown at right. Nucleus outlines were determined using the expression of the HVC and RA marker *PVALB*, not shown. Sections are coronal, scale bar is 100 $\mu$m.

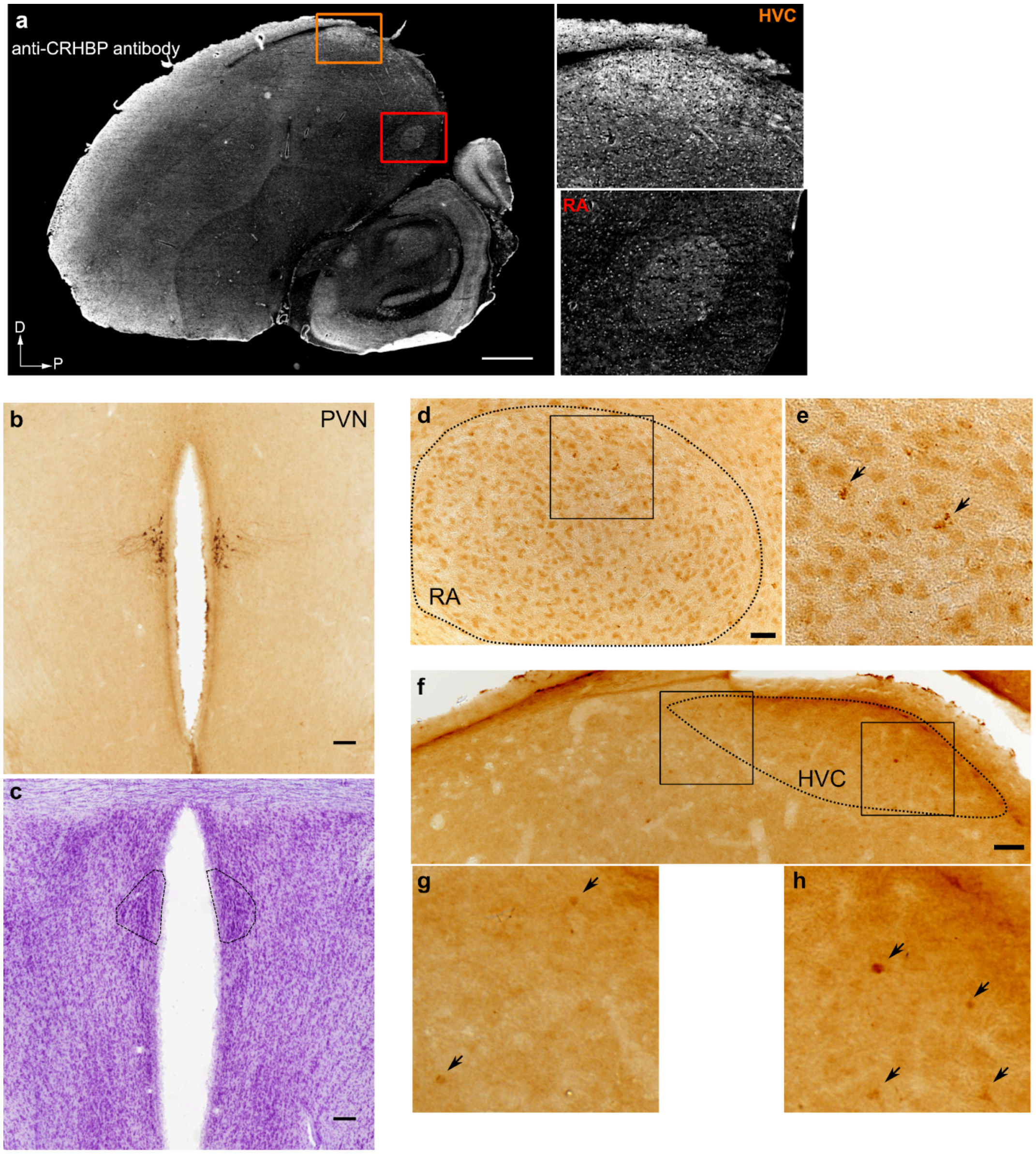


**Supplemental Figure 7. Validation of CRHBP and CRH protein expression in the song motor pathway**
**(a)** Immunofluorescent assay of CRHBP expression across a sagittal section of an adult male Bengalese finch brain. Boxes highlight HVC and RA, which show elevated CRHBP levels. Scale bar is 1 mm. D, dorsal; P, posterior. **(b, c)** CRH immunoreactivity (b) and Nissl stain (c) of the paraventricular nucleus of the hypothalamus (PVN), a region with high CRH expression in mammals (Ketchesin et al., 2017). Scale bar is 50 µm. **(d-h)** CRH immunoreactivity in RA (d-e) and HVC (f-h), with arrowheads indicating immunoreactive cells. e, f, and h are zoomed in views of the boxed regions in d and f. Scale bar is 50 µm.

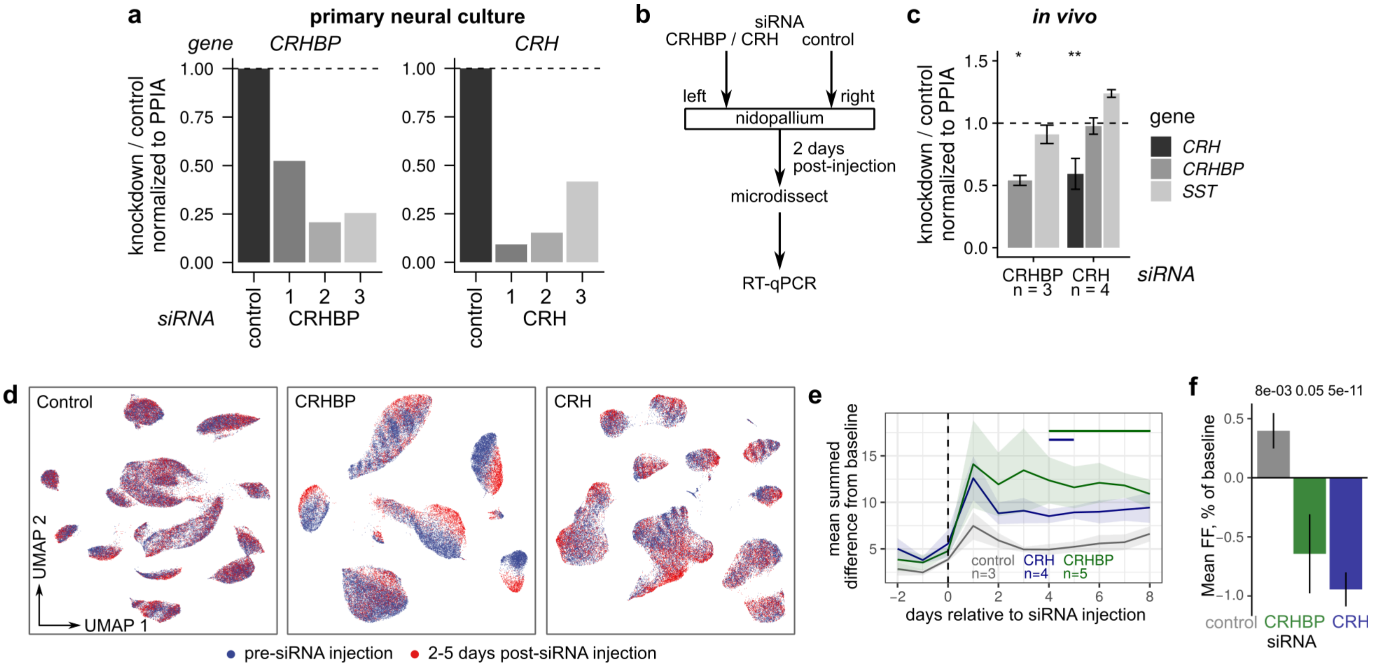


**Supplemental Figure 8. siRNA qPCR validation**

**(a)** *In vitro* validation of siRNA knockdown in Bengalese finch primary neural culture. Three siRNAs were tested for each gene target (CRHBP 1-3 and CRH 1-3). Shown are expression levels for *CRHBP* and *CRH* as assayed by RT-qPCR, normalized by expression of a housekeeping gene (*PPIA*) then by expression after knockdown with a control siRNA. **(b)** Schematic of *in vivo* validation of BrainFectIN knockdown. **(c)** RT-qPCR of *CRHBP*, *CRH,* and *SST* in tissue that had been injected with either control, *CRH*, or *CRHBP* siRNAs. Values are normalized by expression of a housekeeping gene (*PPIA*) then by expression after knockdown with a control siRNA. P-values were calculated using Student's t-test. * $p < 0.05$, ** $p < .01$. **(d)** UMAP plots of songs from three birds with injections of control, CRHBP, or CRH siRNAs in RA. Points are colored by period relative to injection. **(e)** Mean summed differences of UMAP densities for each day relative to pre-injection UMAP densities. Error bands are standard errors of the mean. Color bars indicate days in which values from CRHBP or CRH knockdowns were significantly different from those in control experiments (Student's t-test, two-sided, $p<0.05$). **(f)** Influence of CRH pathway knockdowns on fundamental frequency (FF) mean. For each bird, syllable, and period (pre vs. post-knockdown), the FF mean was computed. Values were then normalized within each bird and syllable to give a percent change relative to pre-knockdown values. These normalized values were then averaged across birds and syllables. Shown here are the normalized post-knockdown values. Pre-knockdown data includes at least 2 days of song prior to injection. Post-knockdown data includes days 2-7 following injection. Error bars are standard errors across birds. A linear mixed-effects model (see Methods) was fit to the data to obtain p-values.
